# Interhemispheric Connectivity Supports Load-Dependent Working Memory Maintenance for Complex Visual Stimuli

**DOI:** 10.1101/2021.03.24.436845

**Authors:** Chelsea Reichert Plaska, Jefferson Ortega, Bernard A. Gomes, Timothy M. Ellmore

## Abstract

A critical manipulation used to study the neural basis of working memory (WM) is to vary the information load at encoding followed by measurements of activity and connectivity during maintenance in the subsequent delay period. The hallmark finding is that delay period activity and connectivity increases between frontal and parietal brain regions as load is increased. Most WM studies, however, employ simple stimuli (e.g., simple shapes or letters) during encoding and utilize unfilled intervals (e.g., a blank screen or fixation cross) during the delays. In the present study, we asked how delay period activity and connectivity change during low and high load maintenance of complex stimuli. Twenty-two participants completed a modified Sternberg WM task with two or five naturalistic scenes as stimuli while scalp EEG was recorded. In each trial, the delay interval was filled with phase scrambled scenes to provide a visual perceptual control with color and spatial frequency similar to the non-scrambled scenes presented during encoding. The results showed that theta and alpha delay activity amplitude was reduced during high compared to low WM load across frontal, central, and parietal sources. Functional connectivity during the delay was assessed by phase-locking value (PLV) and revealed a network with higher connectivity during low WM load consisting of increased PLV between 1) left frontal and right posterior temporal sources in the theta and alpha bands, 2) right anterior temporal and left central sources in the alpha and lower beta bands, and 3) left anterior temporal and posterior temporal sources in the theta, alpha, and lower beta bands. These findings demonstrate a role for interhemispheric connectivity during WM maintenance of complex stimuli. We discuss significance with respect to allocation of limited attentional resources and the filtering of interference.

## Introduction

Previous research supports that the maintenance of information in working memory (WM) depends on activity within a frontal parietal network and includes findings from both fMRI (Bollinger, Rubens, Zanto, & Gazzaley, 2010; Hampson, Driesen, Skudlarski, Gore, & Constable, 2006) and EEG (Babiloni et al., 2004; J. M. Palva, Monto, Kulashekhar, & Palva, 2010). Early direct neural recordings in non-human primates found activity that was sustained throughout the WM delay period (Constantinidis et al., 2018; Kamiński & Rutishauser, 2020; Sreenivasan, Curtis, & D’Esposito, 2014). The persistent activity pattern has been confirmed to some extent in humans, but more recent work demonstrates that activity patterns can be transient (Ellmore, Ng, & Reichert, 2017; Plaska, Ng, & Ellmore, 2021) depending on the length and duration of the delay period, the cognitive strategy employed during maintenance, as well as the type of signal analysis (e.g., time-frequency decomposition, averaging of evoked responses, etc.) used to quantify activity. An even more recent view proposes an activity-silent account of WM in which rapid changes in synaptic weights during the encoding of stimuli support maintenance making it possible to hold online unattended stimuli during a delay period without the need for sustained or transient activity (Beukers, Buschman, Cohen, & Norman, 2021; Kamiński & Rutishauser, 2020; Stokes, 2015). In the activity silent view, the critical neural change supporting maintenance would be a change connectivity not necessarily activity (Babiloni et al., 2004; Gazzaley, Rissman, & D’esposito, 2004; Hampson et al., 2006) induced by altered synaptic strengths.

While empirical findings and theory support both activity and connectivity as neural candidates for WM maintenance, a critical gap in knowledge exists since most studies of WM utilize simple stimuli at encoding (e.g., shapes or letters) followed by unfilled delay periods (blank screens with center fixation crosshairs). Historically, simple stimuli have been used to reduce trial-to-trial variability and to control for novelty. But there are clear downsides to using simple stimuli, which include that they are highly familiar and not ecologically representative. The use of complex stimuli increases the amount of information that must be maintained in WM (Alvarez & Cavanagh, 2004; Awh, Barton, & Vogel, 2007), which likely recruits brain regions in addition to the fronto-parietal circuit typically recruited during maintenance. Complex stimuli like natural scenes place greater demands on attention than simple shapes because they have more features to be combined for perception (Lavie, Hirst, De Fockert, & Viding, 2004). Complexity also requires more executive resources to maintain stimuli as well as filter irrelevant information that could interfere with maintenance (Lavie, 1995). These processes place more demands on prefrontal brain regions and frontoparietal networks (Gruber & Goschke, 2004). Complexity also affects the ability to recode stimuli and use rehearsal as a maintenance strategy (A. Baddeley, 2010, 2012; A. D. Baddeley & Hitch, 1974). Stimuli that are easy to name recruit left-hemisphere speech areas (Gruber, 2001; Smith & Jonides, 1998), while stimuli that are more difficult to name may cause subjects to rely more on right-hemisphere gist-based strategies like attentional refreshing (Cowan et al., 2005).

The use of unfilled delay periods has historically predominated in WM studies because they allow for a clear distinction of delay related activity from background activity that is uncontaminated by perceptual or distractor noise. Neural activity during the WM delay period, when task-related information is being maintained after stimuli have disappeared, is called delay activity (Sreenivasan & D’Esposito, 2019). In the absence of any stimuli during the delay period, attention is assumed to be fully engaged and is thought to be reflected by sustained activity in prefrontal and parietal cortex in both animals and humans (Constantinidis et al., 2018; Goldman-Rakic, 1995; Sreenivasan et al., 2014). Increased alpha power and event-related synchronization measured with EEG have also been identified as correlates of WM maintenance during delay periods in posterior temporal-parietal (Feredoes, Heinen, Weiskopf, Ruff, & Driver, 2011; Sarnthein, Petsche, Rappelsberger, Shaw, & Von Stein, 1998; Scheeringa et al., 2009) and superior parietal regions (Jensen, Gelfand, Kounios, & Lisman, 2002; J. M. Palva et al., 2010), which likely reflects suppression of potentially interfering neural processes (Jensen & Mazaheri, 2010) to minimize external distraction (Bonnefond & Jensen, 2012). In the absence of visual stimuli during the delay period, increases in theta (Jensen & Tesche, 2002) and beta band activity (Jensen & Mazaheri, 2010) in frontal regions have been observed, potentially reflecting engagement of attentional processes and control during stimulus maintenance. In addition to increased activity during the delay period, increased connectivity in the lower frequency bands including theta and alpha have been attributed to attentional processes (J. M. Palva et al., 2010; Zhang, Zhao, Bai, & Tian, 2016), while higher bands like beta and gamma have been attributed to stimulus representation (J. M. Palva et al., 2010). Increases in right hemisphere frontoparietal connectivity have been linked to sustained attention during WM (Coull, Frith, Frackowiak, & Grasby, 1996), while connectivity between frontal-visual regions and frontal-posterior temporal regions has been implicated in supporting maintenance and executive control over maintenance (Daume, Gruber, Engel, & Friese, 2017; Gruber & Goschke, 2004; Rezayat et al., 2021).

Unfilled delay periods have low ecological validity because rarely do we perform day-to-day tasks requiring WM in the absence of any concurrent stimuli. Attention plays a different role at each stage of WM, including selective attention directed externally during encoding and focused attention directed inward during maintenance (Gazzaley & Nobre, 2012). Encoded information captures attention and is prioritized over any distracting information that is to be ignored (Gazzaley & Nobre, 2012). However, research has demonstrated that distracting information can automatically capture attention along with task-related information (Lavie & Cox, 1997). Examining distraction during WM can therefore provide insight into how attention supports maintenance of relevant information while filtering out irrelevant information, a process linked to changes in functional connectivity between prefrontal and posterior brain areas (Gazzaley & Nobre, 2012).

Two different experimental manipulations are commonly employed to study the role of attention during WM. One involves the presentation of task-irrelevant information that is intended to be ignored (Cowan, 2011). The other involves the addition of a secondary task (Morey & Cowan, 2005; Ricker, Cowan, & Morey, 2010) that is not related to the primary task goal. Both methods of external interference can negatively impact performance (Chen & Cowan, 2009; Clapp, Rubens, & Gazzaley, 2010). The impact of distractors is related to the attentional demands of the task, which for WM is manipulated by increasing either the number or complexity of stimuli to be maintained (Alvarez & Cavanagh, 2004; Chen & Cowan, 2009; Simon, Tusch, Holcomb, & Daffner, 2016; Xu & Chun, 2006). As task demands increase, all attentional resources available are used automatically in the early stages processing stimuli, according to the load theory of attention (Lavie, 1995). Once the perceptual load (i.e., the amount of perceptual processing required for a stimulus) has reached capacity, there should be no resources available to process other stimuli that may be presented, including distractors (Lavie, 1995; Lavie & Cox, 1997; Simon et al., 2016). Research shows that increased task demands, including increased WM load, results in reduced ability to process distractors (Sörqvist, Dahlström, Karlsson, & Rönnberg, 2016). Reduced distractor processing is likely to occur during a dual-task requirement or when distractors are unexpected (SanMiguel, Corral, & Escera, 2008). Studies have found that distraction influences connectivity in attention networks that support encoding and maintenance (Greene & Soto, 2014). Interference during maintenance has also been found to disrupt the connectivity between the prefrontal regions and visual regions (Yoon, Curtis, & D’Esposito, 2006) as well as prefrontal and parietal regions (Greene & Soto, 2014). These long-range connections are important for the top-down control of WM (Gazzaley & Nobre, 2012; Von Stein & Sarnthein, 2000) and allocation of attentional resources (Sauseng, Klimesch, Schabus, & Doppelmayr, 2005), processes that are particularly important with increased task demands during WM including maintenance of higher loads. Theta, alpha, and beta frequency oscillations all show increases during the delay period with increasing WM load (Jensen & Tesche, 2002; Mazaheri & Jensen, 2010; Pavlov & Kotchoubey, 2020; Scheeringa et al., 2009; Tuladhar et al., 2007). Gamma band increases have also been implicated during the delay period (Khursheed et al., 2011) and especially during encoding (Mainy et al., 2007; Pavlov & Kotchoubey, 2020). While increased delay activity in lower frequency bands is associated with increased WM load (see (Pavlov & Kotchoubey, 2020), changes in functional connectivity, a method to estimate the degree of interaction among brain regions (Fell & Axmacher, 2011; Fries, 2005), has also been reported during WM maintenance (Payne & Kounios, 2009; Tóth et al., 2012; Zakrzewska & Brzezicka, 2014; Zhang et al., 2016).

Taken together, there is a clear need to understand how neural activity and connectivity changes support the maintenance of complex stimuli at increasing WM load during delay periods filled with perceptually similar distractors. The goal of the present study was to identify the neural and connectivity patterns that support WM maintenance for complex visual stimuli. Participants maintained either a low or high load of complex stimuli during the delay period while looking at phase-scrambled stimuli that lacked semantic content but contained similar spatial frequency and color. The scrambled stimuli during the delay therefore served as a perceptual baseline for the unscrambled scene stimuli presented during encoding. Participants were not required to remember the scrambled stimuli presented during the delay, but a separate behavioral study that we conducted showed that their presence during the delay impacted subsequent memory, suggesting that they induced visual interference. This allowed us to address with EEG measurements how activity and connectivity change as function of WM load in the presence of visual distractors during the delay period. Attention is presumed to be engaged throughout the delay period and attentional mechanisms are often implicated in changes in delay activity (e.g., increased alpha activity as reported by Jensen and Mazaheri (2010)). This raises two questions. What happens when the demand on attention is increased with complex visual stimuli and increasing load? How does activity and connectivity change when the delay period contains interfering stimuli? Additionally, while many EEG delay activity and connectivity studies focus exclusively on frontoparietal interactions (Sauseng et al., 2005), analysis of widespread brain connectivity is critical for understanding how complex visual stimuli are maintained because processing of complex stimuli may involve additional maintenance processes including associations with semantic knowledge (Von Stein & Sarnthein, 2000) that simple stimuli may require less of or not at all. This additional processing may be reflected by more temporal and inter-hemispheric processing.

The hypothesis in the present study is that introducing perceptually similar visual information during the delay period will engage attention and distract from maintenance. We predict that if delay activity is a correlate of maintenance and attention is crucial for successful maintenance, the introduction of interference will have a negative impact on performance and be reflected in changes in delay activity and connectivity. When there is a greater demand on attentional resources, such as during maintenance at higher WM load, there are fewer resources available to process the interfering stimuli. Therefore, in a low-load WM condition that requires fewer attentional resources, there will be more resources available to deal with interfering stimuli presented during the delay. As a result, there will be reduced delay activity during high load WM, especially in frontal regions critical for attention and maintenance and performance will be negatively impacted. Additionally, it is predicted that there will be reduced frontoparietal connectivity during the delay period to support filtering of interference. At low WM load, the interference will have less of an impact on the delay activity and connectivity. Performance will therefore be better than the high load condition, and delay activity and connectivity in frontal and parietal regions will be greater for the low compared to high load condition.

## Methods

### Participants

The study protocol was approved by the Institutional Review Board of The City University of New York Human Research Protection Program (CUNY HRPP IRB). All methods were carried out in accordance with the relevant guidelines and regulations of the CUNY HRPP IRB. Because the data were collected as part of a simultaneous EEG/MRI study, all participants completed an MRI safety questionnaire. After review and approval by the MRI director, participants provided informed consent after the study procedures were described to them verbally. The study consisted of 24 subjects but two subjects were excluded from all analyses because of unusable EEG data or failure to complete both task conditions (e.g., because they fell asleep during one of the task conditions or misunderstood task instructions) which left a final sample of 22 subjects (12 females, 18 to 54 years old, mean age 24.95 years, SD = 8.57).

### Memory Tasks

This study employed a variant of the classic Sternberg WM task (Sternberg, 1966) with low (two stimuli) and high (five stimuli) loads. Participants completed 50 trials of each load. The loads were presented in randomized order (i.e., participant 1:low-load first, high-load second; participant 2: high-load first, low-load second, etc.). Approximately ten minutes later, participants completed an immediate recognition task. Before going into the MRI scanner, participants completed three practice trials for each WM condition and the recognition task inside the MRI control room before being placed inside the scanner. Before completing each task inside the scanner, participants were read the same instructions that were given to them during the practice trials.

Each WM task condition consisted of an encoding phase (with two or five stimuli each presented for 1400 ms each, *Figure 1*), a delay period (6000 ms), a yes/no 50/50 probe choice (1400 ms) followed by jitter (∼3000 ms) consisting of a blank screen. During the delay period, six phase-scrambled scenes were presented each for 1000 ms. Phase scrambling preserves color and spatial frequency information but removes semantic content, which allows for them to serve both as a perceptual baseline relative to the encoding condition and as distractor stimuli relative to a static blank screen with fixation cross (see *Supplemental Material*). Participants were instructed to look at but not remember the phase-scrambled stimuli during the delay while maintaining the scene stimuli presented to them during the encoding phase. The evidence that these phase scrambled stimuli served as interference during the delay is demonstrated by their detrimental effects on memory found in a separate behavioral study that we conducted before this EEG study (see *Supplemental Material*). For the probe choice task, participants were presented with either a new scene (negative probe) or a previously presented old scene (positive probe) after which they had to signal by button press if they saw the scene before or not. Participants responded using a fiber optic response pad (fORP, Current Designs, Inc., Philadelphia PA) that they held in their right hand. They pressed a green button on the left of the pad for “Yes” and a red button on the right for “No”.

**Figure 1.**
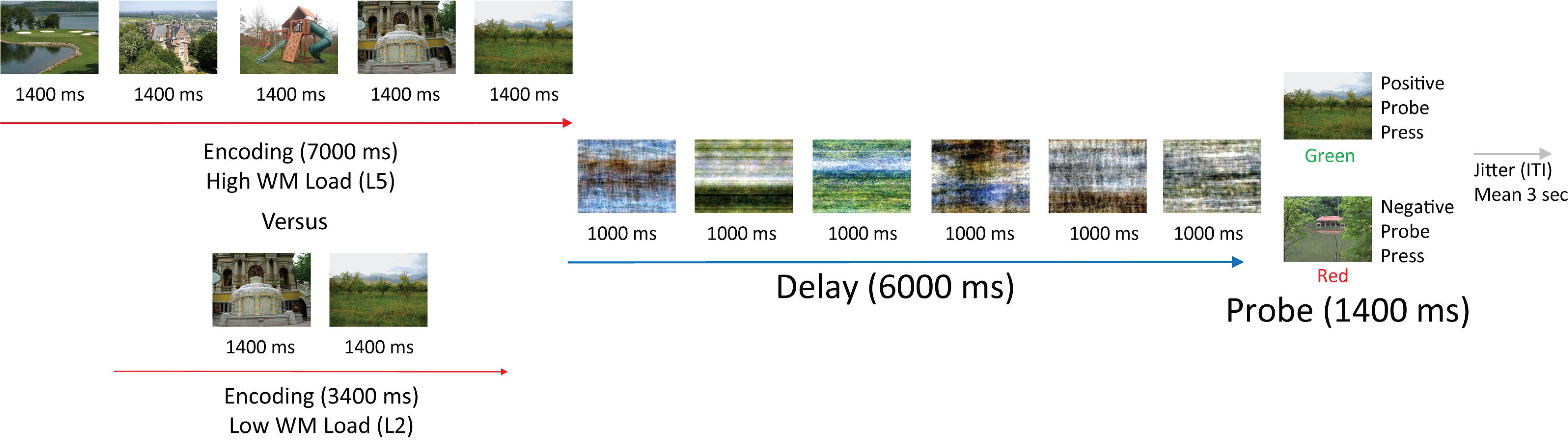
High- and Low-load Working Memory Task Trial Layout. The task consisted of two working memory loads: a low load (two scene stimuli) and a high load (five stimuli). In the high load condition (top): participants were presented with five images in succession (1.4 sec each), a delay period with six phase-scrambled images (1 sec per image for a total delay period duration of 6 sec), a probe choice (1.4 sec), which was either one of the earlier presented stimuli or a new image, and a jitter period (∼3 sec) with a blank gray screen that indicates the end of the trial. For the low-load condition (bottom): participants were presented with two images in succession (1.4 sec each), a delay period with 6 phase-scrambled images (1 sec per image for a delay period total of 6 sec), a probe choice (1.4 sec), which is either one of the earlier presented images or a new image, and a jitter period (∼3 sec) with a blank gray screen that signaled the end of the trial. Participants completed both loads in randomized order.

After completing the WM tasks, participants completed an immediate recognition task, which consisted of new and old scenes (from the L2 and L5 WM condition) intermixed with phase-scrambled scenes. During presentation of the phase-scrambled scenes, participants were instructed to look at them and not to make a response. For all other stimuli, participants were told to indicate if the scene was new (i.e., never seen before) or old (i.e., from one of the two earlier WM tasks). The tasks runs were interspersed with five-minute rest conditions in which participants fixated a black cross while instructed to stay awake and keep their eyes open. Compliance was monitored using an in-bore eye-tracking camera. The tasks took approximately 45 minutes to complete, and the total time within the scanner was approximately 90 minutes.

The scene stimuli were color outdoor naturalistic scenes from the SUN database (Xiao, Hays, Ehinger, Oliva, & Torralba, 2010) sized at 600 by 800 pixels and displayed on a gray background. Stimuli were randomly presented within each trial using an BOLDscreen LCD monitor that was located behind the MRI bore. Participants viewed the monitor in a mirror above them attached to the head coil. Before beginning the task, the experimenters confirmed that participants were able to see the monitor clearly. During the experimental session, participants were not able to see the button box. They were asked to memorize the location of the green and red buttons and were asked to press the red and green buttons separately to confirm they know the correct yes/no button mapping before beginning the task.

### EEG Acquisition

We collected continuous 32-channel EEG from passive electrodes (31 scalp electrodes and one ECG electrode) inserted in Brain Vision MR-safe caps while simultaneously acquiring structural and functional MRI. One subject had EEG recorded from 64 channels (63 scalp and one ECG electrode), but for comparison with the other participants a 32-channel montage was applied to process this participant’s data. This subject was included in all EEG analyses except for the connectivity analysis described below because a custom source montage was not available. Scalp electrodes were arranged according to the 10-10 international system. In accordance with the Brain Products simultaneous EEG-fMRI acquisition guidelines (Brain Products BrainAmp MR Operating and Reference manual Version 4.0), *AgAL* MR-safe gel was used to bring all electrode impedances to 20 kOhm or below. An ECG electrode was placed on the back left shoulder blade to record ballistocardiogram. EEG caps were prepared and fitted outside the scanner and impedances were checked again after positioning on the scanner bed and movement into the bore. For safety, impedances were monitored throughout the scanning session. If impedances exceeded 35 kOhm between scans, the participant was removed from the bore, and more gel was applied until the impedance was lowered. After needed corrections, the participant was repositioned and returned to the bore and impedances were checked again before repeating a localizer scan and continuing with recording.

EEG data were recorded at 2500 Hz. Each participant was positioned 40 mm caudal to isocenter in the bore of the 3.0 Tesla Siemens scanner (Mullinger, Yan, & Bowtell, 2011). Initially, the first few subjects (n = 5) were recorded at 500 Hz or 1000 Hz and sample rate was increased to 2500 Hz to improve fMRI artifact cleaning. Changes in sampling rate were based on manufacturer recommendations during experimental start-up. After fMRI artifact cleaning was completed, data from all participants were down-sampled to 500 Hz.

### EEG Processing

EEG processing was done using BESA Research 7.0. MRI artifact correction was carried out using the BESA Research fMRI artifact gradient removal algorithm with the Allen Method (Allen, Josephs, & Turner, 2000) following the default BESA settings (number of artifact occurrence averages = 16). This method includes a sliding-window template with 8 artifact occurrences before and after the artifact that are used to create the artifact template, which is then subtracted from the data. The duration of each functional MRI volume was automatically detected (repetition time = 2000 msec) and was based on the user-defined fMRI trigger code (R128) recorded during the session, which appeared after the initial three dummy scans. For the first 19 participants, the fMRI trigger code was not recorded in the EEG files and was instead substituted by a trigger code recorded in the file (Sync-On timestamped every 2000 ms). To align the fMRI gradient artifact with the alternate trigger, the first trigger for each file was manually adjusted in the MRI Artifact Removal settings as the delay between marker volumes and the start of volume acquisition.

Ballistocardiogram (BCG) correction was carried out using the ECG electrode channel. For BCG detection, a low cutoff filter of 1 Hz (zero phase, 12dB/oct) and a high cutoff filter of 20 Hz (zero phase, 24 dB/oct) were applied as recommended by BESA (BESA Research WIKI, 2020). A template BCG cycle was manually selected for each participant in each file by a trained research assistant. A pattern-matching algorithm was then used to identify the PCA components that explained the BCG pattern (usually between 4-5 components depending on variance accounted for). Blink correction was carried out using a similar method (Ille, Berg, & Scherg, 2002; Picton et al., 2000). The method of correction removes the variance associated with a blink (or BCG pattern), from each channel, using the template pattern selected. Blink correction was carried out using the frontal electrodes Fp1 and Fp2 since the EEG-cap did not contain designated electrooculogram channels. Prior to beginning the WM and recognition tasks, participants were asked to blink five times in a row for the purposes of creating a blink template. If the cued blink artifact resembled a spontaneous blink, this blink artifact was input to the pattern matching algorithm to identify the PCA component. Otherwise a more representative natural blink was selected. If the PCA components accounted for greater than 97% of the variance, it was accepted. Otherwise the pattern-matching algorithm was run again with another template blink. Artifact-corrected data were used for all analyses to ensure that the BCG artifact would not distort the findings. The reference electrode was the frontal pole (Fpz) during recording and was re-referenced offline to the common average reference for the initial sensor-level analyses.

### Time Frequency Analyses

There were two WM conditions, a low- (two scene stimuli) and a high-load (five scenes) each with the same delay period length (6000 ms). All analyses described below compared the low- to high-load delay period conditions (L2 vs. L5). Individual trials were excluded based on BESA Research criteria for artifact rejection. Trials were excluded for amplitude > 150 μV, gradient > 75 μV, low-signal criteria > 0.1. After rejection, the average number of usable trials was 40 for L2 and 42 for L5 out of 50 total trials per condition. There was no difference between the number of useable trials between conditions (p = 0.32). Time-frequency analysis was carried out at both the sensor- and source-levels. Complex demodulation of the recorded EEG signals for each trial was done in BESA Research v7.0 (Hoechstetter et al., 2004; Papp & Ktonas, 1977). A detailed description of the demodulation can be found in Ellmore et al. (2017). The timeframe (t) under consideration was the full delay period (0 to 6000 ms) and the baseline was average amplitude across the full epoch (Ellmore et al., 2017). The selected frequency (f) ranges were 4 to 40 Hz: theta (4-8 Hz), alpha (8-13 Hz), and beta (13-30 Hz), and lower gamma (30-40 Hz). Beta is further subdivided into lower (13-20 Hz) and upper beta (20-30 Hz). Time-frequency matrices were generated using a finite impulse response (FIR) filter with latencies of 100 ms and 0.5 Hz frequency steps. The amplitude within each frequency and latency bin was generated for each WM load (Time Frequency Amplitude or TFA). TFA is expressed as the absolute value of the amplitude in microvolts (µV). The amplitude within a frequency and time bin is expressed relative to the amplitude of the baseline (Temporal Spectral Analysis or TSA). The TSA is expressed as either a positive or negative percent change in amplitude with the equation:

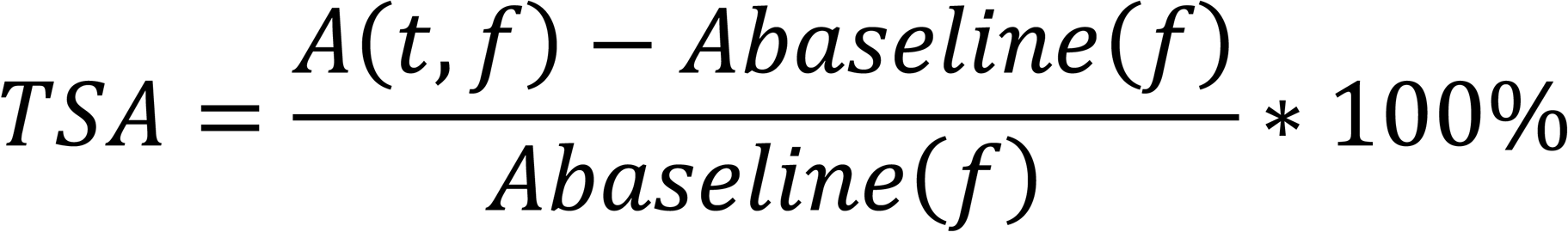

where,

*A(t,f) = amplitude during the timeframe of interest and frequency*

and

*A_baseline_ (f) = mean amplitude in the frequency band during the baseline period*

If TSA is positive, it reflects synchronization of activity relative to the baseline period. If it is negative, it reflects desynchronization (Pfurtscheller, 2001; Pfurtscheller & Da Silva, 1999).

To estimate source-level delay period activity, the BESA default Brain Regions montage was applied to the delay period TSA to account for the potentially overlapping sources of delay activity (Ellmore et al., 2017). The Brain Regions montage is comprised of 15 discrete regions including frontal, temporal, and parietal regions (see *Supplemental Figure 2 and Supplemental Table 2*). The source-level montage estimates brain region sources by reducing the overlapping signals from the scalp electrodes by calculating weighted combinations of the recorded signal (Hoechstetter et al., 2004; Scherg, 1992). The spatial separation of these fixed sources ensures there is minimal cross-talk between the regions (Scherg, Berg, Nakasato, & Beniczky, 2019).

### Connectivity and Phase-Locking Analyses

To quantify interactions between brain regions during the delay period, phase-locking value (PLV) analyses were run on the time-frequency data. Both coherence and PLV measure the amount of oscillatory synchronization between two brain regions (Lowet, Roberts, Bonizzi, Karel, & De Weerd, 2016). Coherence is a measure of the linear covariance between two signals in a particular time-frequency bin, taking into account both phase and amplitude of the signal under the assumption that the signal is stationary in time (Rosenberg, Amjad, Breeze, Brillinger, & Halliday, 1989). Phase-locking value is a measure of the phase of signal, without respect to the amplitude for a specific frequency-time bin, which is then compared with the phase of another signal within the same frequency-time bin (Lachaux, Rodriguez, Martinerie, & Varela, 1999). While coherence and phase-locking values may overlap in the connections that are identified, phase-locking value has no assumptions about linearity and stationarity and so it is considered a more robust measure of oscillatory synchronization (Lowet et al., 2016). Moreover, synchrony between two regions only needs to consider the phase of two signals to make determinations about communication (Lachaux et al., 1999). Therefore, to make more accurate assumptions about long-range and cross-hemispheric differences, only phase-locking values (PLV) were compared.

In the PLV analysis, the Brain Regions source montage was applied to the data to reduce the number of comparisons (i.e., sources are fewer than sensors). The time-frequency analysis was run on the delay period using the same parameters described above. TSA was computed with complex demodulation between 4-40 Hz with 0.5 Hz/100 sec steps, to be consistent with the above analysis. Phase-locking values were computed in BESA Connectivity v1.0 for both the low- and high-load conditions. For PLV, the values range from 0 (no synchrony) to 1 (completely synchronous) and values between 0 and 1 represent partial synchrony (Lachaux et al., 1999). The equation for PLV is:

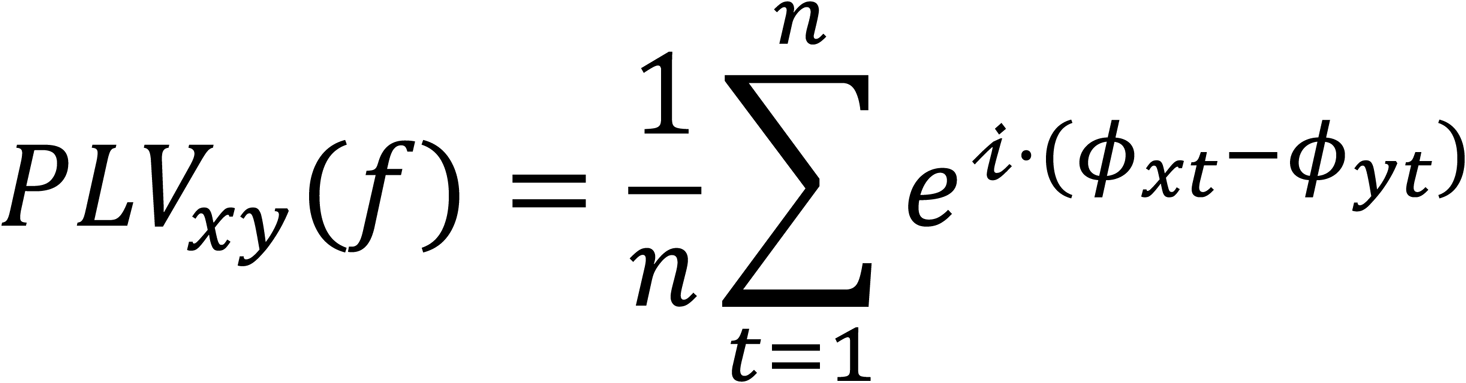

where,

f=frequency of interest, n=number of time points (t) in the epoch, i=imaginary number

and

ϕ_x_ and ϕ_x_= phase angles from two signals x and y with the frequency of interest

Spurious results outside of the selected frequency ranges were ignored (BESA Connectivity, 2020). BESA Connectivity outputs were converted to a BESA Statistics compatible format in Matlab v2020a using BESA generated scripts. 3D head plots depicting changes in connectivity between brain regions were generated using BrainNet Viewer (Xia, Wang, & He, 2013).

### Statistical Analyses

The behavioral performance on the WM and recognition tasks was analyzed using custom Matlab (v2020a) scripts. Performance is reported as the percent correct out of the total possible trials. Reaction time is reported as the mean reaction time for responses across all trials. The kurtosis and skewness of performance and reaction time was evaluated in SPSS v24.0. Non-parametric statistics using the related samples Wilcoxon Signed Rank Test were computed after reviewing the skewness and kurtosis of each dependent variable. Since the behavioral distributions indicated moderate skewness and kurtosis, a non-parametric paired-samples test was used for the performance comparisons. Three subjects who produced many no response trials on either the low- or high-load condition were initially included in the behavioral analysis and subsequent EEG comparisons. These subjects were excluded from the correct trials only analysis (see *Supplemental Material*), and the behavioral analysis was repeated excluding these subjects. Excluding these three subjects did not change the behavioral findings. The results reported below also exclude them. Regression and pointplots of behavioral performance and mean delay activity were run in Python v3.6 using Seaborn v0.11.1 and can be found in *Supplemental Material*.

All delay activity analyses were carried out in BESA Statistics v 2.0, which deals with the multiple comparisons problem by carrying out permutation tests producing cluster and associated probability values (Maris & Oostenveld, 2007). Paired t-tests were used for the delay activity comparisons across conditions, followed by correlations with behavioral measures. The cluster value is a sum of t-values for a t-test and a sum of r-values for correlation across a group of adjacent bins. A bin consists of sensors that are <4 cm distance, latency of 100 msec, and frequency bins of 0.5 Hz. A null distribution is created by sampling randomly from clusters across subjects and across time-frequency bins. Significant clusters are summed t- or r-values within a specific time-frequency bin that exceeds a specific threshold which are then compared to the random null distribution created from 1000 permutations (Bullmore et al., 1999; Ernst, 2004; Freedman & Lane, 1983; Maris & Oostenveld, 2007; O’Gorman, 2012). Clusters were considered significant if the p-value was less than or equal to .05. A more detailed description of the theory behind and computation of the EEG permutation tests can be found in Ellmore et al. (2017). For visualization, significant clusters are indicated with a mask that is colored either blue or orange. Orange represents low-greater than high-load, and blue represents high-greater than low-load for the TFA and TSA (event-related synchronization) comparisons. For connectivity, significant clusters are indicated with either red representing greater phase-locking for low-compared to high-load or blue representing greater phase locking for high-compared to low-load.

## Results

### Behavioral

There was a significant difference between low- and high-load WM task performance (Figure 2, low-load 94.74%, SD 6.37, high-load 86.21%, SD 10.24, related samples Wilcoxon signed rank test, p = .001). There was also a significant difference between low- and high-load on the immediate recognition test (low-load 70.38%, SD 14.84, high-load 61.43% correct, SD 17.62, related samples Wilcoxon signed rank test, p = .002). Reaction times for low-load WM trials were faster than for high-load (low-load 878.62 ms, SD 139.76, high-load 955.71 ms, SD 122.02, related samples Wilcoxon signed rank test, p = .002).

**Figure 2.**
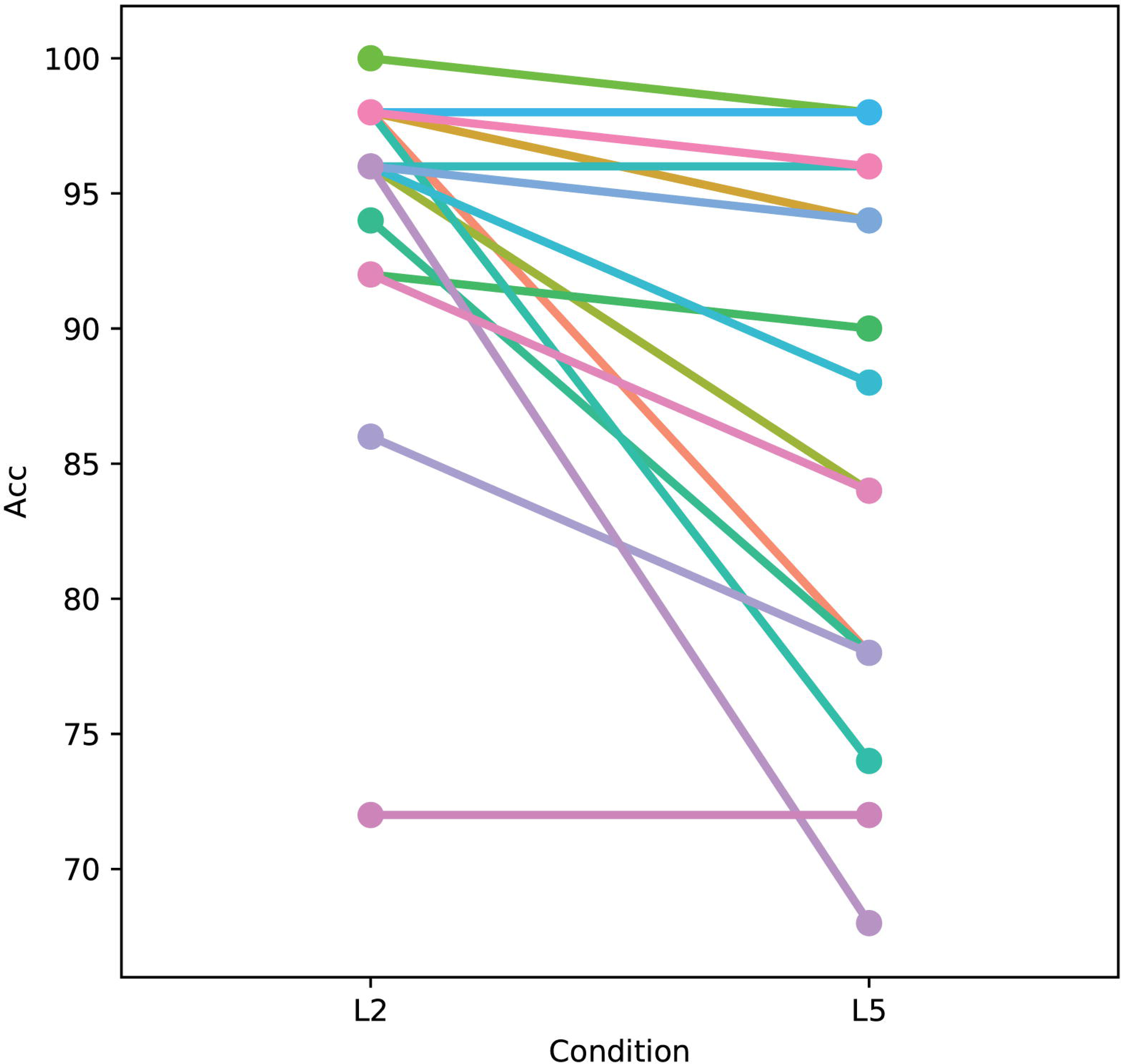
Better Performance on the Scene Working Memory Task for the Low-than the High-Load Condition. Pointplot of performance on the low- (L2) and high-load (L5) WM tasks. Each line in the figure represents a subject. Performance was significantly better on the low-compared to the high-load condition (94.6% correct vs. 86.2%, Wilcoxon Signed Rank Test p = .001). Each color represents an individual subject. Accuracy (Acc) was defined as percentage of trials correct out of total possible trials. Three subjects with a large number of no response trials were excluded.

### Correlations with Behavior

Correlations between performance and delay activity showed no significant correlations between TSA delay activity and performance for either condition (low-load p-value = 0.09; high-load p-value = 0.32). There were also no significant correlations between the phase-locking values and performance for either condition (low-load p-value = 0.80; high-load p-value = 0.63).

### Absolute Amplitude

A sensor-level comparison of low- and high-load delay period absolute amplitude revealed five significantly different clusters of delay activity (*Table 1*). Four of the five clusters revealed greater amplitude for the low-compared to high-load condition throughout the entire delay period (0-6000 ms) spanning all frequencies. The clusters were mainly in the alpha and beta ranges, with greater amplitude for the low-compared to high-load condition (Figure 3a-c). The remaining cluster showed a greater amplitude for the high-load condition in the first half of the delay period (0-2600 ms). This cluster encompassed right centro-frontal channels (F4’_avr, C4’_avr, FC6’_avr, and CP6’_avr) and included the upper beta and gamma range (frequency = 29-37 Hz). This analysis was conducted on all delay period trials, regardless of correct or incorrect probe response. To investigate whether activity during incorrect responses may have influenced this difference, a separate analysis examined the relationship between activity in selected significant clusters with performance (*Supplemental Figure 2a-d*) but included only delay periods for trials with correct responses. The findings with respect to absolute amplitude remained unchanged (see *Supplemental Material*).

**Figure 3.**
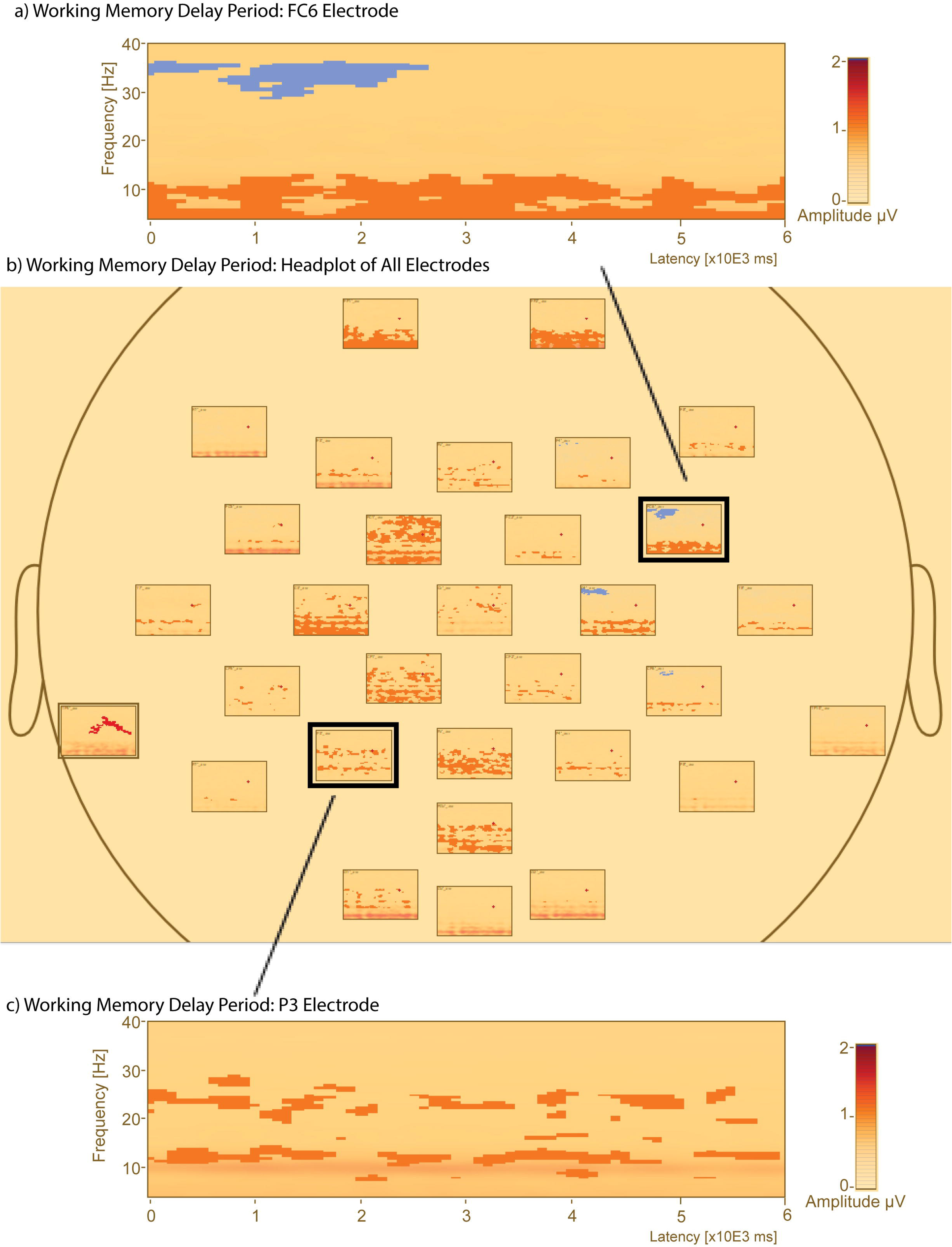
Comparison of Absolute Amplitude Delay Period Activity in Low- and High-Load Conditions Reveals Greater Alpha and Beta Band Amplitude Across the Delay Period. Select absolute amplitude plots in the left frontal and right parietal regions of the delay period revealed five clusters of significant differences in activity (*p* < .05). Orange clusters represent low-load delay activity greater than high-load while blue clusters represent high-load delay activity greater than low-load. The y-axis shows frequency (Hz); x-axis shows the time in sec. a) Time Frequency Absolute Amplitude plot for the FC6 electrode. The plot shows the low-load condition Absolute Amplitude with the significant clusters overlaid as a mask. b) Head plot of the overall pattern of absolute amplitude difference during the delay period for all sensors. c) Time Frequency Absolute Amplitude plot for the P3 electrode. The plot shows the low-load condition Absolute Amplitude with the significant clusters overlaid as a mask. The selected FC6 (a) and P3 (c) electrodes are highlighted by a dark box on the head plot (b).

**Table 1:**
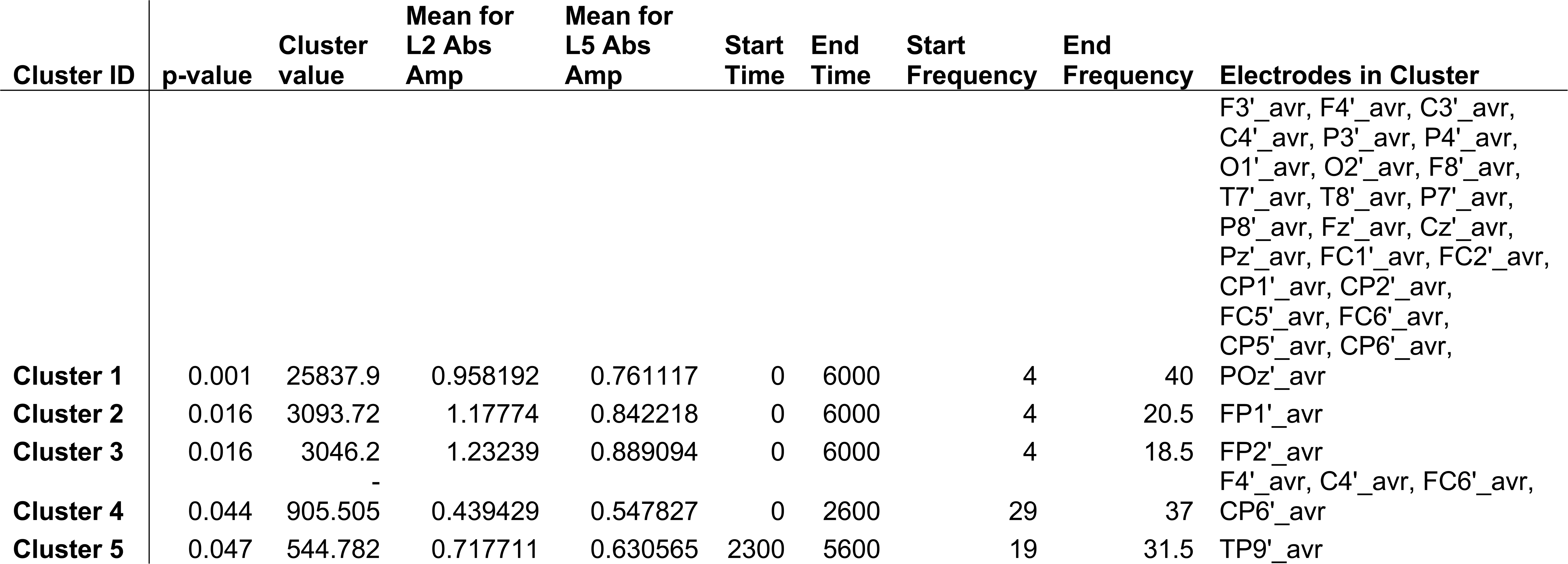
Significant clusters of absolute amplitude difference between the low- (L2) and high-load (L5) conditions during the delay period (Time: 0 to 6000 msec) for all sensors.

### Temporal Spectral Amplitude

A sensor-level comparison of low- and high-load delay period temporal spectral amplitude revealed a transient pattern of delay activity that was most pronounced in the right parietal region (Figure 4a-b, P8). Early event-related synchrony (ERS) was found in the alpha and lower beta band beginning about 1000 ms into the delay and lasting until approximately 2000 ms. Event-related desynchrony (ERD) was found in alpha, lower beta, and upper beta beginning about 3500 ms into the delay period and continuing to the end. The pattern of ERS and ERD was more pronounced in the low- compared to high-load condition but was not significantly different. There was only one cluster of significantly different TSA delay activity. It was found early in the delay period (time = 300-2500 ms) where TSA during low-load exhibited greater ERD than high-load (Figure 4b-c, cluster value = -1337.84, p = 0.013) in the upper beta and gamma range (frequency = 29.5 – 40 Hz). The significant difference across the delay period encompassed 26 of the 31 electrodes with a right lateralized focus.

**Figure 4.**
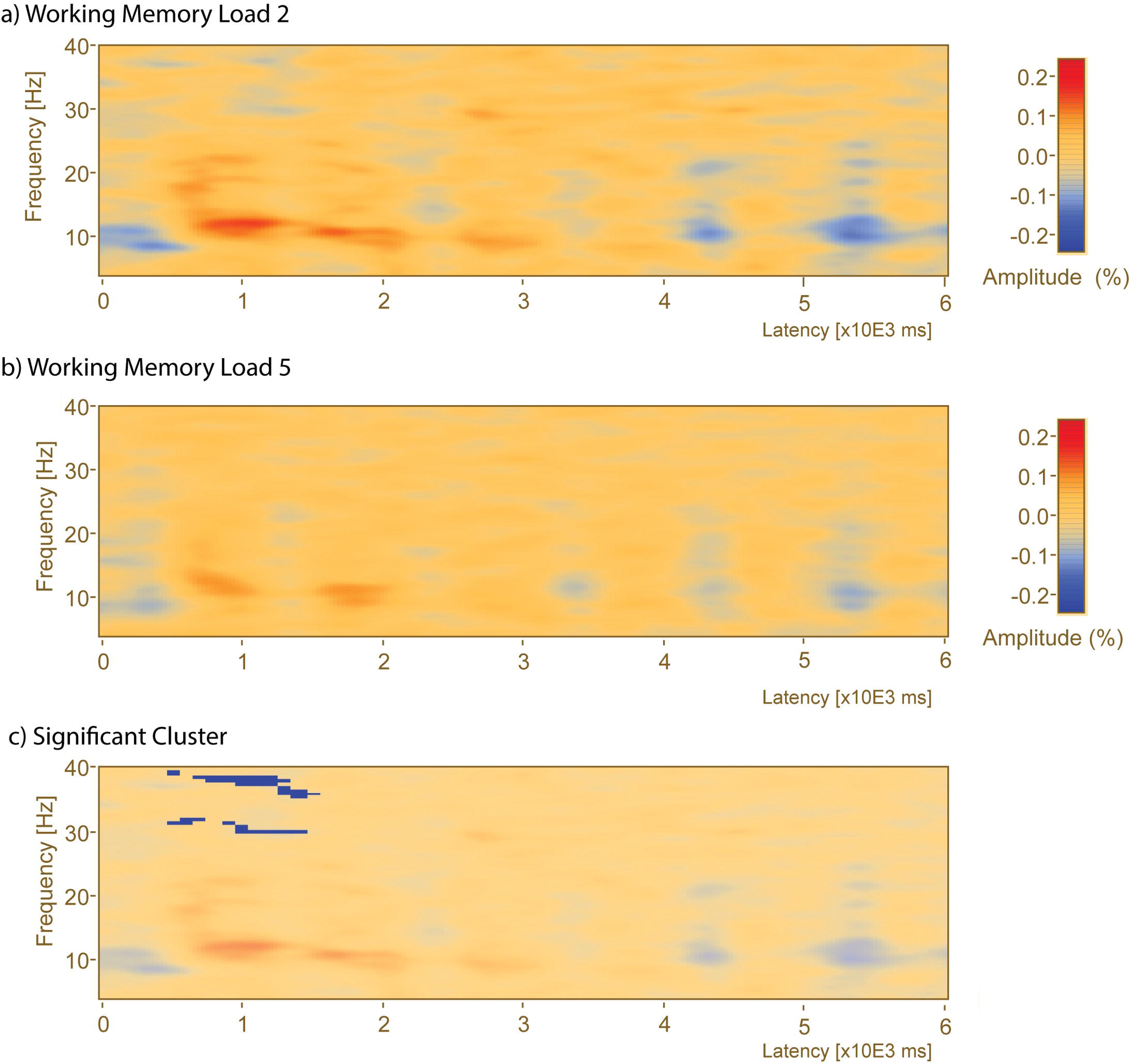
Comparison of Delay Period Time Frequency Between Low- and High-Load Conditions Reveals Early Gamma Band ERD. A whole window time Frequency analysis plot for the P8 electrode selected based on results from previous research in our lab highlighting the importance of the right parieto-occipital region (Ellmore et al., 2017) for this task. a) Time Frequency Analysis for the P8 electrode in the low-load condition. b) Time Frequency Analysis for the P8 electrode in the high-load condition. c) The plot shows the Time Frequency Analysis plot from the low-load condition with the significant load difference cluster overlaid as a mask. The blue cluster represents reduced ERD for the high-load condition. Review of the overall pattern of delay activity between the low- and high-loads reveals a similar transient pattern of delay activity for the parieto-occipital censors with an early period of increased synchronous activity in the upper alpha and beta bands (500 msec to 3000 msec) followed by a period of desynchronous activity in the same frequency bands (4000 msec to 6000 msec).

### Temporal Spectral Amplitude Source Analysis

A brain regions source-level comparison of low- and high-load delay period temporal spectral amplitude found two clusters of significantly different delay activity (Figure 5). The left frontal region exhibited greater event-related synchronization in the low-compared to high-load condition during the first half of the delay period (Cluster 1: cluster value = 176.60, p = 0.003, frequency = 4-8.5 Hz, ERS greater during low load, time = 2700-3300 ms at source FL_BR). The left temporoparietal region exhibited greater event-related desynchronization for the low-compared to high-load condition (Cluster 2: cluster value = -134.03, p = 0.036, frequency = 22.5-27 Hz, ERD greater for low-load, time= 1800-2600 ms at source TPL_BR) in the middle of the delay period.

**Figure 5.**
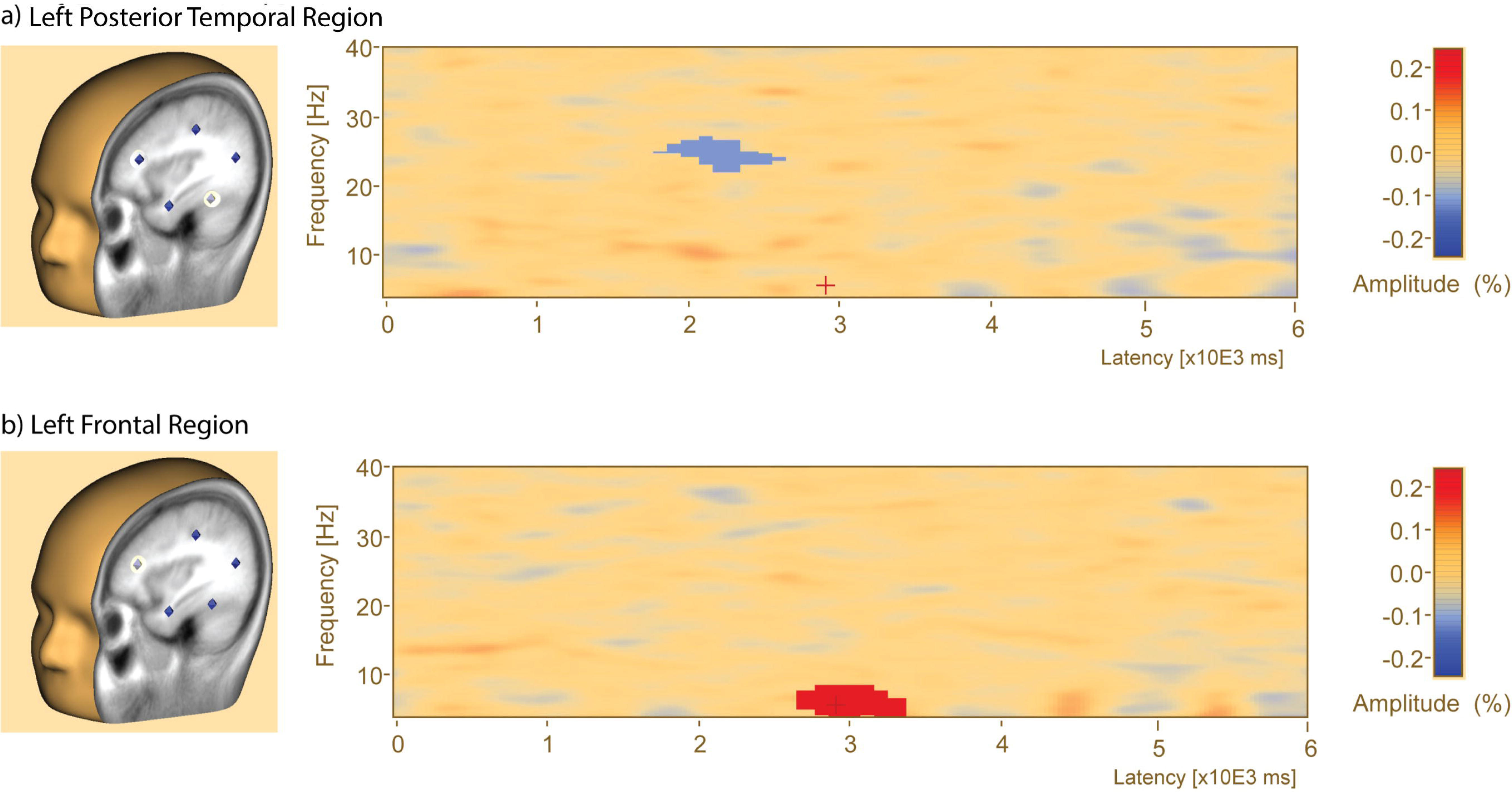
Delay Period Activity Source Analysis Reveals Changes in Left Posterior Temporal and Left Frontal Regions During Maintenance. Time Frequency Analysis plots during the delay period during the brain regions source analysis. a) 3-D view of Brain Regions Montage superimposed on an MRI structural scan with the Left Posterior Temporal Region highlighted in yellow (left). Time Frequency Analysis for the Left Posterior Temporal Region during the delay period (right). The blue cluster represents greater ERD in the upper beta range for the low-load condition. b) 3-D view of Brain Regions Montage superimposed on a template MRI structural scan with the Left Frontal Region highlighted in yellow (left). Time Frequency Analysis for the Left Frontal during the delay period (right). The red cluster represents greater ERS in the theta and alpha range for the low-load condition.

### Phase Locking Value Source Analysis

Phase-locking value connectivity for both the low- and high-load conditions appeared sustained throughout the entire delay period (*Figure 6*). A paired-samples t-test with corrections for multiple comparisons revealed six significantly different clusters between low- and high-load (*Table 2*). The matrix for both loads showed increased left intra-hemispheric connectivity among anterior temporal, temporoparietal, and parietal regions across all frequency bands. There was inter-hemispheric connectivity but it was restricted to the lower frequency bands including increased connectivity between midline and bilateral central regions and frontal regions in the upper beta and gamma bands. There was also increased connectivity between frontal polar midline and bilateral parietal regions in the alpha band and between occipital polar midline and bilateral parietal regions in the gamma and alpha bands.

**Figure 6.**
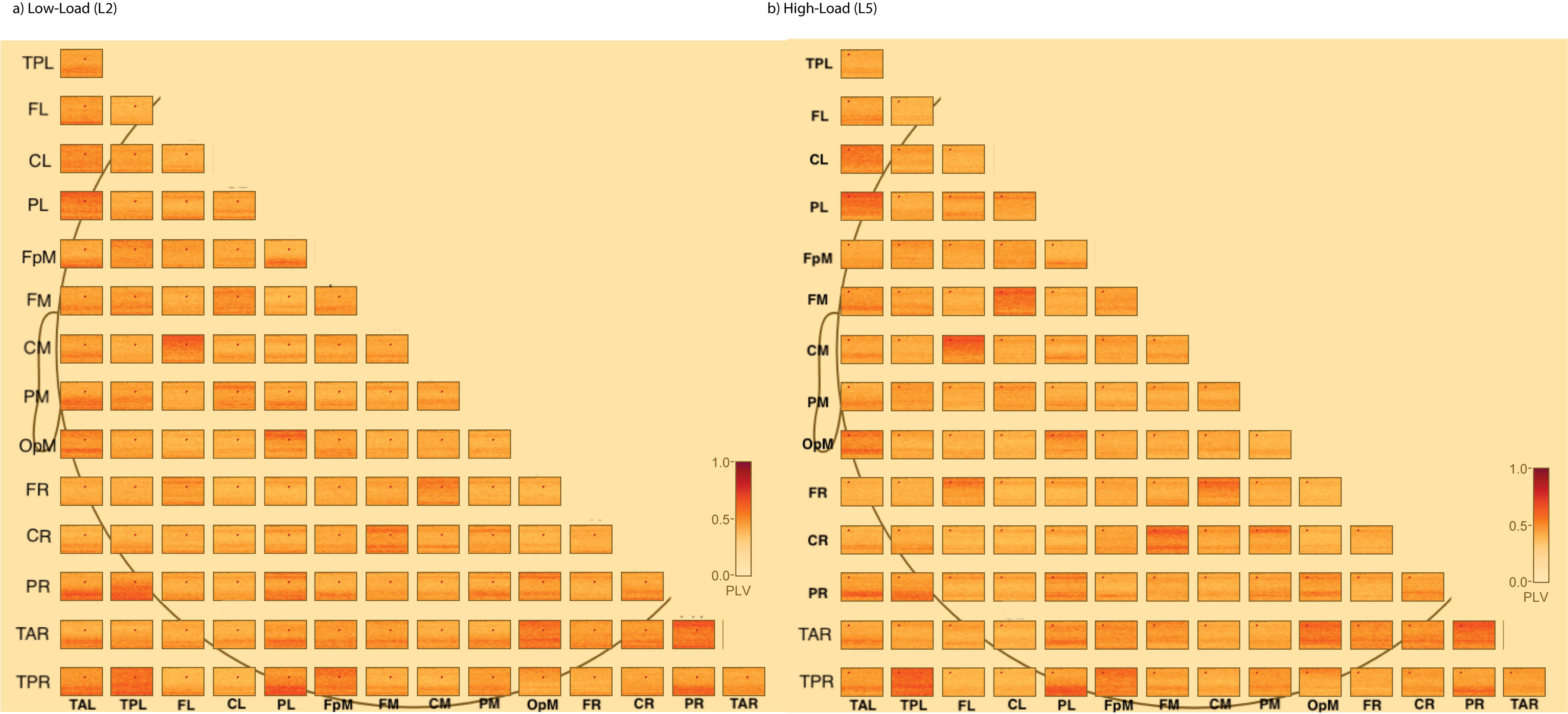
Phase Locking Value Connectivity Matrices for the Low- and High-Load Conditions. Each subplot represents a matrix of connections among 15 brain regions (see Methods) within a condition, averaged across trials and subjects. For each matrix, the major X and Y axis contain labels for the brain regions. Each box inside the matrix represents 1 connection (e.g., top left box – connection between left posterior temporal – left anterior temporal). For each box, the plot shows the phase-locking value (PLV) for that connection and displays PLV on a scale from 0 – 1, with 0 (yellow color) indicating no connectivity and 1 (deep red) indicating highest level of connectivity. For each box, the Y axis is frequency from 4-40 Hz and the X axis represents the full delay period (0-6000 msec). a) Connectivity matrix for the low-load condition (L2). b) Connectivity matrix for the high-load condition (L5). Both the low- and high-load conditions appear to have sustained levels of PLV throughout the entire delay period (see *Brain Regions-Phase Locking Value* section).

**Table 2:**
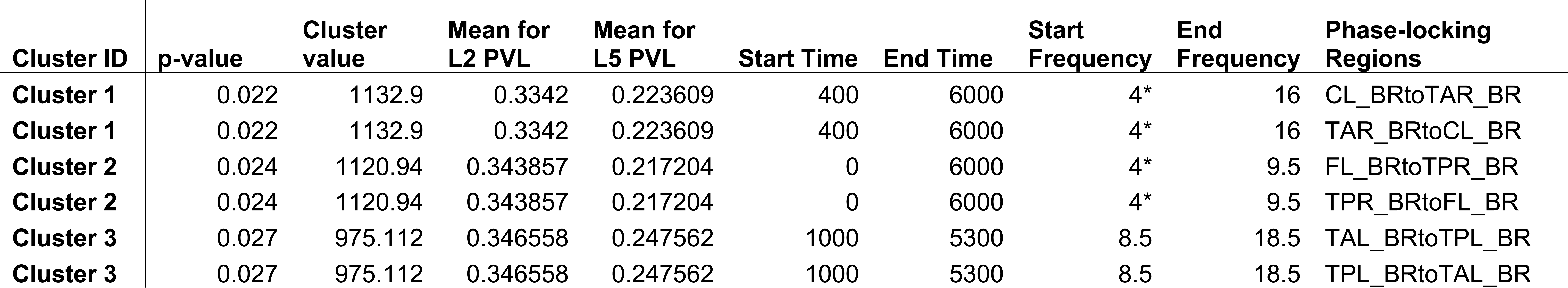
Significant clusters of phase-locking value (PLV) between 15-brain regions during the delay period (Time: 0 to 6000 msec). Each PLV connection between the low- (L2) and high-load (L5) conditions has two significant clusters associated with it to represent the opposite connection (e.g., PL – FL and FL – PL). The PLV results do not provide information about the directionality. The start frequency for all comparisons was 4 Hz, but can erroneously report frequencies lower (i.e., lower than 4 Hz), so start indicates numbers that were adjusted due to erroneous values (see *Methods*).

There were three clusters of significantly different PLV including a cluster between left frontal and right posterior temporal sources (Figure 7a, cluster value = 1120.94, p = 0.024, frequency range 4-9.5 Hz, time range 0-6000 ms, sources FL_BR - TPR_BR). There was also greater PLV between left frontal and right posterior temporal regions in the theta and alpha ranges throughout the entire delay period. There was a significant difference in PLV between left anterior temporal and left posterior temporal (Figure 7b, cluster value = 975.11, p = 0.027, frequency range 8.5-18 Hz, time range 1000-5300 ms, sources TAL_BR - TPL_BR). There was significantly greater PLV between left anterior temporal and left posterior temporal regions in the alpha and lower beta ranges throughout the entire delay period. There was a significant difference in PLV between left central and right anterior temporal regions (Figure 7c, cluster value = 1132.9, p = 0.022, frequency range 4-16 Hz, time range 400-6000 ms, sources CL_BR - TAR_BR). There was greater PLV between the left central and right anterior temporal regions in the theta, alpha, and lower beta ranges starting after about 400 ms and continuing until the end of the delay period. The main analysis was conducted on all delay period trials that survived artifact rejection, regardless of correct or incorrect responses. A direct statistical comparison of correct and incorrect trials was not possible due to the low number of incorrect trials. An exploratory analysis was however conducted to examine if incorrect responses may have influenced the results. The relationship between PLV and performance was plotted for significant clusters (*Supplemental Figure 4a-f)*. An analysis conducted using only delay periods on trials with correct responses (see *Supplemental Material*) found that, of the significant clusters listed above, the cluster of significantly different PLV between FL – TPR was no longer significant (*Supplemental Figure 5*).

**Figure 7.**
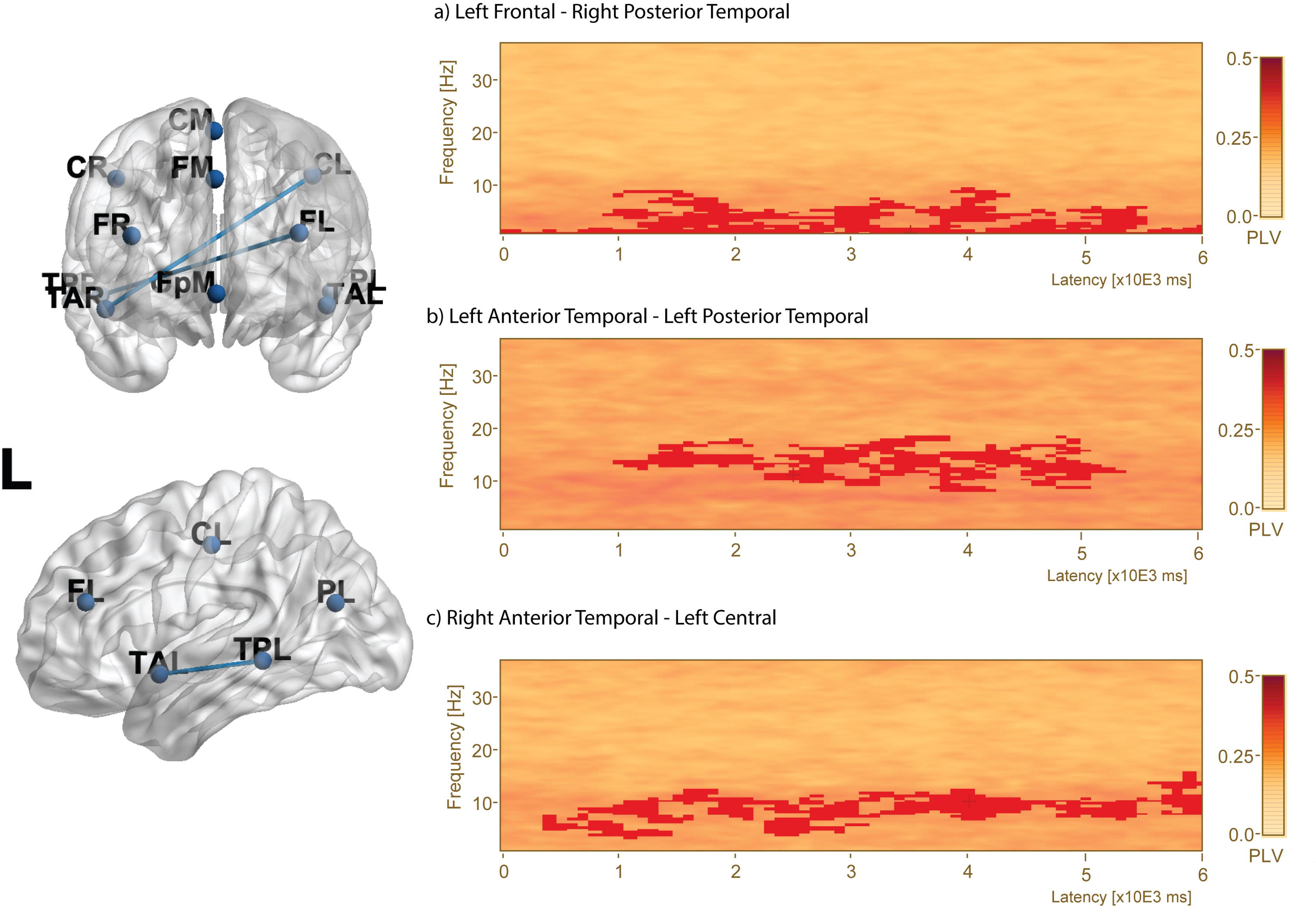
Phase Locking Values During the Delay Period Reveal Three Different Connections with Higher Connectivity During the Low-Load Condition. The plots show phase locking values (PLV) during the delay period in a brain regions connectivity analysis. A 3D transparent brain frontal (left-top) and left lateral view (left-bottom) displaying the significantly different PLV connections highlighted with a blue connection line. Letters on the 3D brain letters refer to locations: L = Left Hemisphere, R = Right Hemisphere, A = anterior, P = posterior, T = Temporal Lobe, P = Parietal Lobe, and F = Frontal Lobe. a) Connectivity plot for the connection between left frontal and right posterior temporal regions with mask of the significant cluster indicating that the low-load condition had greater connectivity between these two regions than the high-load condition (red mask, p = 0.024). b) Connectivity plot of the connection between the left anterior temporal and left posterior temporal regions indicating that the low-load condition had greater connectivity than the high-load condition (red mask, p = 0.027). c) Connectivity plot of the connection between right anterior temporal and the left central regions indicating that the low-load condition had greater connectivity between these two regions than the high-load condition (red mask, p = 0.022).

The other PLV connections remained unchanged when considering correct trials only, and there were three additional clusters identified (*Supplemental Figure 6*).

## Discussion

In the present study, we tested the hypothesis that introducing perceptually similar visual information during the delay period would engage attention and distract from maintenance in a load-dependent manner. We found results consistent with this hypothesis with performance being better in a low-load working memory condition as well on an immediate recognition task for stimuli from the low-load condition. Moreover, the present results provide some support for the persistent delay activity hypothesis as delay amplitude was above baseline throughout the delay for both conditions. Persistent or transient activity is opposite to what would be expected according to an activity silent view of WM where activity would return to baseline for unattended stimuli during the delay period (Stokes, 2015; Stokes, Muhle-Karbe, & Myers, 2020). However, it is impossible to draw firm conclusions about activity-silent accounts based on results from the present study because activity-silent mechanisms may carry information from stimuli encoded during the previous trial, while activity-based mechanisms dominate for the current trial (Barbosa et al., 2020), which was the focus of the current investigation.

We also predicted that there would be reduced frontoparietal connectivity during the delay period to support filtering of interference for phase-scrambled stimuli presented during the delay. At low WM load, this interference was predicted to have less of an impact on delay activity and connectivity. Thus, performance would be better than the high load condition, and delay activity and connectivity in frontal and parietal regions would be greater for the low-compared to the high-load condition. We found patterns in our results that are consistent with these predictions. There was greater amplitude in alpha and lower beta bands for the low-load condition across most sensors. Greater amplitude in the alpha and lower beta range in the low-load condition could reflect greater disruption of attention for the high-load condition by the interfering stimuli (Bonnefond & Jensen, 2012). It may also reflect that during low-load WM maintenance there were more resources available to process the interfering stimuli (Yoon et al., 2006) and the available resources could have been used to deal with the interference. While greater activation of the parietal region during the high-load condition would be consistent with traditional accounts of WM processing scaling positively with the amount of information being maintained, lower activity may instead reflect a different neural mechanism to deal with interference during the delay. The greater perceptual load during the high-load condition may have resulted in a reduced distractor effect on the delay activity (Lavie et al., 2004), while the greater amplitude in the low-load condition could reflect maintenance of both the target and interfering stimuli.

At the sensor-level, transient delay activity was found during both the low- and high-load WM conditions. There was an early pattern of event-related synchronization (ERS) in the alpha and low beta bands until about 2000 ms followed by a pattern of event-related desynchronization (ERD) in the alpha, lower and upper beta bands that began in the middle of the delay period and lasted to the end. Increases in amplitude and ERS in the alpha band are associated with inhibition of brain regions to suppress irrelevant information during maintenance (Jensen & Mazaheri, 2010; Klimesch, 2012). These changes may also reflect engagement of semantic systems that when processing stimuli are then inhibited during maintenance to prevent interference (Klimesch, 2012). Similar transient patterns were found during low- and high-load conditions except for one cluster of activity early in the delay period that showed greater ERD during low- compared to high-load in the upper beta and gamma regions across most sensors. During a source analysis of TSA, differences were found in a left-frontal region in the theta band and very low alpha bands and right-posterior temporal regions in the upper beta band during the early-middle delay period. Event-related desynchronization has also been found by others in the alpha band during maintenance (Koshy et al., 2020). This finding has been attributed to the anticipation of an upcoming response as well as attentional processes (Klimesch, Sauseng, & Hanslmayr, 2007). ERD may reflect semantic access to support the maintenance of the complex stimuli (Klimesch, 2012). Lower beta band ERD has also been associated with semantic processing (Hanslmayr, Spitzer, & Bäuml, 2009). The alpha and low beta band activity may reflect complementary processes (Pavlov & Kotchoubey, 2020). The ERD observed in the latter half of the delay period of the present study is consistent with preparation for an upcoming probe response during an extended period of maintenance (Plaska et al., 2021). As the same type of stimuli were used in both WM load conditions, the pattern of delay activity found during the relatively long six seconds of maintenance could reflect similar processes across both WM loads.

There was greater amplitude in higher beta and gamma bands for the high-compared to the low-load condition in right fronto-central sensors during the first half of the delay period. There was also an early period of higher upper beta and gamma band amplitude and ERS in the same regions. Higher beta and gamma activity have been associated with working memory maintenance (Axmacher et al., 2007; Lundqvist, Herman, Warden, Brincat, & Miller, 2017; Lundqvist et al., 2016; S. Palva, Kulashekhar, Hämäläinen, & Palva, 2011). In addition to being associated with maintenance, higher beta activity has been associated with volitional control (Lundqvist et al., 2017) and may be related to the selection of which information is maintained (de Vries, Savran, van Driel, & Olivers, 2019; de Vries, Slagter, & Olivers, 2020). The application of volitional control may help explain why performance during the high-load condition was far above chance. While competition between the target and the interfering stimuli is possible, the pattern may also represent competition among the five stimuli during maintenance.

### Connectivity and Maintenance

Differences in connectivity between the low- and high-load conditions were found with greater connectivity during the delay period between the left central and right anterior temporal regions in an analysis of all trials and in a separate analysis of only trials with correct responses. The central region is a brain region located superiorly near the junction of the frontal and parietal lobes, potentially overlapping with the anterior portion of the posterior parietal cortex (Whitlock, 2017). The posterior parietal cortex (PPC) is involved in attentional processes (Hutchinson, Uncapher, & Wagner, 2009) with more dorsal regions associated with allocating and control of attention. Increased connectivity among occipital, temporal, and parietal regions including PPC has been reported with increased WM load. Specifically, increases in lower frequency bands, such as alpha and lower beta, are attributed to the deployment of attentional resources (J. M. Palva et al., 2010; S. Palva et al., 2011). Increased connectivity between left central and right anterior temporal regions within these lower frequency bands may reflect control of attention for the maintained stimuli, which was likely easier at a low load of two stimuli compared to five stimuli, and which is important for preventing the degradation of the WM representation (Lorenc, Mallett, & Lewis-Peacock, 2021).

There was also greater connectivity between left frontal and right posterior temporal regions, and between the left anterior temporal and the left posterior temporal regions. Left frontal regions are important for filtering interference and increased attentional selection (Gazzaley & Nobre, 2012; Jha, Fabian, & Aguirre, 2004). The right posterior temporal regions include the temporoparietal junction, which is implicated in the control of attention (Geng & Vossel, 2013). This region is also implicated in the storage of the encoded stimuli (Feredoes et al., 2011; Scheeringa et al., 2009) and selection of stimuli into WM (Rezayat et al., 2021). If the posterior temporal region encompasses the superior temporal gyrus, this pattern could be consistent with storage of the encoded stimuli as Park et al. (2011) proposed that the superior temporal gyrus represents a store for complex visual stimuli. While filtering is often associated with frontoparietal connectivity, the connectivity pattern has typically been demonstrated using simple stimuli. Filtering associated with complex stimuli like those used in present experiment would be essential for correct responses. Consistent with this idea, significant differences in this network emerged only when all trials were included in the analysis and was no longer present when examining only correct trials. If this network supports the filtering of interfering stimuli during the maintenance of complex stimuli, then the observation that there was no significant difference between the low- and high-load delay periods (e.g., when examining delay periods on correct only trials) suggests that filtering was critical for successful performance. Therefore, including incorrect trials in the analysis resulted in diminished connectivity within this network in the high-load condition consistent with an inability to filter the interference, particularly during incorrect trials.

There was also an increase in short-range connectivity between the left anterior temporal region and left temporoparietal region. The left anterior temporal region is critical for language, short-term memory, and semantic associations (Boucher, Dagenais, Bouthillier, Nguyen, & Rouleau, 2015; Helmstaedter, Elger, Hufnagel, Zentner, & Schramm, 1996; Hermann, Wyler, Bush, & Tabatabai, 1992). The left posterior temporal region is a critical junction for somatosensory information and has been implicated in word processing and comprehension (Binder et al., 1997). Phase-locking between these two regions was restricted to the alpha and lower beta regions, which suggests that this connection represents feedback from the anterior temporal to the posterior region throughout the latter half of the delay period (about 1000-5000 ms). This may represent a process in which the encoded stimulus is recoded into a label using an association of stored semantic knowledge followed by rehearsal. Even without explicit instructions to engage in rehearsal, participants could adopt a dual-coding strategy involving verbal recoding and rehearsal, particularly with stimuli that contain high semantic content (Brown, Forbes, & McConnell, 2006) like the naturalistic scenes used in the present study. Increased connectivity was observed during the low-load condition, which is consistent with behavioral observations and subject self-reports that it is easier to rehearse two stimuli with recoded with semantic associations than it is to rehearse five stimuli. In the high-load condition, participants may have had to rely on an alternate maintenance strategy (Chen & Cowan, 2009) such as less efficient gist-based attentional refreshing, which may explain load-dependent differences in performance.

### Attention and Maintenance

Maintenance of complex stimuli, including naturalistic scenes, likely requires an increased need for attentional resources (Chen & Cowan, 2009). There may have been no attentional resources remaining to process the interfering stimuli, regardless of load, since resources are automatically captured by complex scenes (Lavie, 1995; Lavie & Cox, 1997; Simon et al., 2016). Complex scenes contain many components, which need to be combined (e.g., a palm tree with an ocean) to produce semantic meaning (e.g., a beach). Participants draw meaningful connections between stimulus features and stored knowledge (Asp, Störmer, & Brady, 2021), which likely requires attentional resources. Even at a low load of only two stimuli, the complexity and associated meaning of the picture may have used up the available resources. Future studies should include comparisons of complex scenes with simple stimuli, such as objects, to confirm if interference differentially impacts maintenance for complex scenes and simple objects during increasing WM load.

External interference has been introduced in WM tasks in multiple forms, including interfering stimuli that a participant is instructed to ignore (Clapp et al., 2010). As in the current task, if the interfering stimuli are from the same category (e.g., using face stimuli with face distractors), disruption of performance is greater than if the distractor is from a different category (Yoon et al., 2006). Interfering stimuli that have similar content, and not just the low-level features of spatial frequency and color in the phase-scrambled stimuli used in the current experiment, are more likely to take attentional resources for processing (Yoon et al., 2006). In fact, research with face stimuli and face distractors compared with phase-scrambled distractors, have found that only the same-category stimuli and not phase-scrambled distractors were disruptive to both performance and delay activity (Dolcos & McCarthy, 2006; Dolcos, Miller, Kragel, Jha, & McCarthy, 2007). It is possible therefore that the phase-scrambled stimuli used in the present experiment did not actually interfere with WM maintenance. A dual-task paradigm may have been better at capturing attention, such as a judgement task (Clapp et al., 2010), a saccade task (Postle & Hamidi, 2007) or verbal retrieval from a pre-learned word list (Ricker et al., 2010). Future research should examine if same-category visual stimuli (e.g., indoor scenes) or a dual-task may disrupt attention differently and impact delay activity.

If increased delay activity and connectivity are expected with increasing load, as has been reported in many delay period studies (J. M. Palva et al., 2010; Zhang et al., 2016), the increased delay activity is presumed to reflect that all items are held in working memory (e.g., with a high WM load of five with all five stimuli maintained simultaneously). The studies that have used high load manipulations tend to use simple stimuli (e.g., colored squares with nameable colors or easy-to-name objects). It is possible that reduced amplitude activity and reduced phase-locking for the high-load condition found in the present experiment could reflect a processing associated with more complex stimuli such as scenes. In the low-load condition, changes in amplitude and synchronization likely reflect the representation of two items, no matter how complex, as holding onto two stimuli is considered well below typical working memory capacity (Alvarez & Cavanagh, 2004). On the other hand, the high-load condition is above or near the maximum number of stimuli that can be maintained, although this capacity may differ based on the individual (Adam, Vogel, & Awh, 2017). As a result, the maintenance period of the high-load condition may be reduced, both in terms of amplitude activity and connectivity, because participants were not able to maintain all stimuli simultaneously. Depending on an individual’s capacity, the maintenance period may only reflect two or three items at a time due to their complexity, despite the instructions to maintain all stimuli. Both the findings of the planned analysis including all trials and the additional analysis using only correct trials, are consistent with this possibility.

It is also possible that maintenance was protected from interference because participants engaged in maintenance rehearsal. Participants may have automatically recoded the visual stimuli into verbal labels and rehearsed them during maintenance, despite no instructions to do so. Studies have found that the automatic recoding of verbalizable stimuli, such as the naturalistic scenes used in this experiment, will occur even without explicit instructions to do so (Postle, D’Esposito, & Corkin, 2005). Rehearsal is not an attention-demanding process that will interfere with maintenance of visual information (Vergauwe, Camos, & Barrouillet, 2014), but linking semantic associations during rehearsal with visual information, which may occur during processing of complex naturalistic scenes (Plaska et al., 2021), may capture attentional resources. If the processing and linking of semantic associations occurs during maintenance, then interference may not have occurred because any available attentional resources would be used up and not available to process the distractors. Recently this claim about rehearsal has been revisited with the idea that rehearsal may actually place a demand on attention (Thalmann, Souza, & Oberauer, 2019), which could also explain why the interfering stimuli did not impact performance especially in the low-load condition, when rehearsal is more likely to have been carried out. Future studies should attempt to dissociate the attentional demands of rehearsal with complex visual stimuli and the filtering of interference.

### Limitations

One obvious limitation of this study is the small sample size (n=22), which is typical of EEG and MRI studies, due to costs and the difficulty in finding suitable participants (Plaska et al., 2021; Rose et al., 2016; Wei & Zhou, 2020). A second limitation is that EEG collected in the supine condition while laying inside a scanner may show different patterns of amplitude, including reduced cortical activity and laterality differences (Spironelli & Angrilli, 2017). However, the overall cortical patterns found in the present study are consistent with EEG in an upright sitting position (Scheeringa et al., 2009). To reduce the amount of time spent during the experiment while keeping the number of trials across conditions consistent, the two WM load conditions contained only 50 trials each. A larger number of working memory trials per condition may have resulted in better signal-to-noise ratio (Luck, 2014) and enabled a formal comparison between correct and incorrect trials. Finally, unlike the behavioral experiment that motivated this study the experimental design of the main EEG experiment did not include a control delay period (e.g., a delay period with blanks screen or simple fixation cross) in which there were no interfering stimuli present (Postle & Hamidi, 2007). Consequently, it could be argued that the differences in performance and delay activity are simply due to the working memory load difference (2 images vs 5 images) and are unrelated to the interfering stimuli that were originally designed to serve as a perceptual baseline. We argue that this is not the case as the observed delay activity for the high-load condition was reduced. At increasing WM load, studies commonly report increased delay activity (Jensen et al., 2002; Jensen & Mazaheri, 2010; Tuladhar et al., 2007). Future studies should include a non-distractor control (i.e., unfilled delay period) to confirm these differences are due to the presence of interfering stimuli.

### Conclusions

The presence of interference during the delay period of a visual working memory task impacted maintenance for complex stimuli in a load-dependent manner. Successful maintenance of complex visual stimuli in the low-load condition as evidenced by better performance, could be the result of attentional filtering of interfering stimuli, the ability to focus attention on maintenance, and the application of verbal maintenance strategies. At the neural level, greater reductions in amplitude were observed in the alpha and beta ranges for the high-load condition suggesting that the presence of interfering visual stimuli during the delay period resulted in a distractor effect. Increased left frontal-right posterior temporal, left anterior temporal-posterior temporal connectivity, and right anterior temporal-left central connectivity during the low-load condition suggests a combination of intra- and inter-hemispheric interactions may explain protection against interference during WM maintenance of complex visual stimuli.

## Acknowledgements

We thank A. Duke Shereen, PhD and Kenneth Ng for their help with task set-up and with data collection. We want to thank Farzana Antara, Gabriela Silva, and Albert Perdomo for help with EEG data processing. We also want to thank Michael Epstein, PhD for his constructive feedback on the manuscript.

## Availability of Data

Deidentified data will be made available upon reasonable request to the corresponding author.

## Author Contributions

T.M.E. conceived the experiment, obtained funding and developed the task. C.R.P., J.O., and B.G. carried out EEG preparation, data collection, and carried out all data processing. C.R.P. carried out all data analysis and wrote the manuscript. J.O. and B.G. reviewed and provided feedback on the manuscript. T.M.E. aided in interpreting the results and writing the manuscript as well as supervised all aspects of the experiment.

