## Supplemental Material for "Interhemispheric Connectivity Supports Load-Dependent Working Memory Maintenance for Complex Visual Stimuli"

### Supplemental Methods

The experimental design of the EEG study (see *Methods*) was motivated by the results of a separate behavioral study. In the behavioral experiment, a different group of participants completed a Sternberg working memory (WM) task with a similar design. It consisted of two conditions, a low-load condition (two stimuli, *Supplemental Figure 1a*) and a high-load WM condition (five stimuli, *Supplemental Figure 1b*). Participants completed trials with both loads in randomized order, followed by an immediate recognition task and a separate long-term recognition task 24 hours later. During the WM trials, participants saw either a standard fixation cross or phase-scrambled scenes during the delay period. An effect of load was found for both the WM task and immediate recognition task. For the long-term recognition task, there was a significant interaction of load and delay period condition with performance that was lower for the high-load condition when the scrambled images were presented during the delay relative to simple fixation during the delay (*Supplemental Figure 1c).* The behavioral difference as a function of delay period condition (scrambled vs. fixation) motivated the present EEG study because it suggested that the presentation of phase-scrambled scenes during the delay period induces more interference during maintenance as evidenced by reduced performance on the subsequent memory task. The Sternberg WM task is an ideal design for studying load manipulations and attention during maintenance because this task requires selective attention to be directed towards stimuli during the various stages of working memory (Gazzaley & Nobre, 2012). The load manipulation ensures that there are different levels of task demands on attention (Lavie, 2005). This task is also ideal for EEG analysis of memory as phases of the task are separated into encoding, maintenance, and retrieval.

### Supplemental Results

The planned load comparisons were initially conducted using both correct and incorrect trials to maximize the signal-to-noise ratio, especially for the high-load condition where there were fewer correct trials. To determine whether delay activity during trials with incorrect responses influenced differences in amplitude and connectivity between conditions, we re-analyzed the data using trials with correct responses only. Studies that have examined delay period activity associated with correct and incorrect probe responses suggest that overall activity is reduced (Vogel & Machizawa, 2004; Zhang, Zhao, Bai, & Tian, 2016) and as a result exclude incorrect trials and missed response trials from their analyses. This correct trial only comparison was run after the planned comparisons using all delay trials were completed and after an update to BESA Statistics was released. The BESA Statistics update v2.1 eliminated the step in which connectivity data is converted to a compatible format in Matlab (BESA Connectivity, 2020), which allowed for this analysis to be completed exclusively in BESA Statistics v2.1.

To examine whether there was a relationship between the absolute amplitude (*Supplemental Figure 3*) and connectivity findings (*Supplemental Figure 4*), we first plotted the mean activity per subject per cluster against performance. A performance effect is not apparent for any of the significant clusters, for either amplitude or phase-locking, but these findings may be influenced by our questionable lowest performers. As described in the methods section, three participants had several no response trials. One participant omitted most responses on the low load condition, while two participants omitted most responses during the high load condition. As we were not able to confirm whether the participants were responding incorrectly but very slowly or having other issues with the button box (in the scanner they cannot see the buttons as they were supine with the button box out of view), we eliminated these participants from the supplemental analysis.

In this supplemental analysis, the average number of trials per condition was 38.05 for the low load condition and 35.47 for the high load condition ($p = 0.21$, n.s.). The reduction in the average number of trials, particularly in the high-load condition is consistent with reduced performance on the high-load condition (see *Behavioral Results*). Significant clusters of absolute amplitude difference between the low- and high-load conditions in this correct only trial analysis are shown in *Supplemental Table 2*. The differences in amplitude were similar to the original analysis (see *Results*) with four significant clusters of absolute amplitude activity greater for the low-load condition across most sensors. There were also two significant clusters of absolute amplitude activity greater for the high-load condition with a right fronto-central focus. The difference in activity expanded slightly to include the right central parietal region as well.

Significant clusters of PLV between the low- and high-load conditions in the correct trials only analysis are listed in *Supplemental Table 3*. The most significant cluster from the analysis including all trials (Cluster 1: CL – TAR) remained. PLV between CL – TAR was greater for the low- compared to high-load condition. Additionally, the cluster representing TAL – TPL remained after this repeated analysis but was only marginally significant (Cluster 9/10 and Cluster 11/12, p = 0.086 and 0.1 respectively). In the analysis with all trials, the PLV was increased for the low-load condition and was sustained throughout most of the delay period. After analysis with only correct trials, the increased PLV for the low-load condition was evident early in the delay period (time: 0-2300 msec) and again later in the delay period (time: 3200-5700). The cluster that did not survive the analysis with only correct trials was the connection between FL – TPR. Review of the raw PLV plots from the low- and high-load conditions suggests that they have similar patterns of increased phase locking throughout the delay (*Supplemental Figure 5*). There were three additional significant clusters of PLV with this repeated analysis (*Supplemental Figure 6*). There was increased PLV for the low-load condition between FpM – PL throughout most of the delay (time: 600-4900 msec) in the alpha and low beta band. There was increased PLV for the low-load condition between OpM – TAR throughout most of the delay (time: 1100-5000 msec) in the theta and alpha band. Finally, there was increased PLV for the low-load condition between TPL – PR early in the delay (time: 0-2100 msec) in the low and upper beta band.

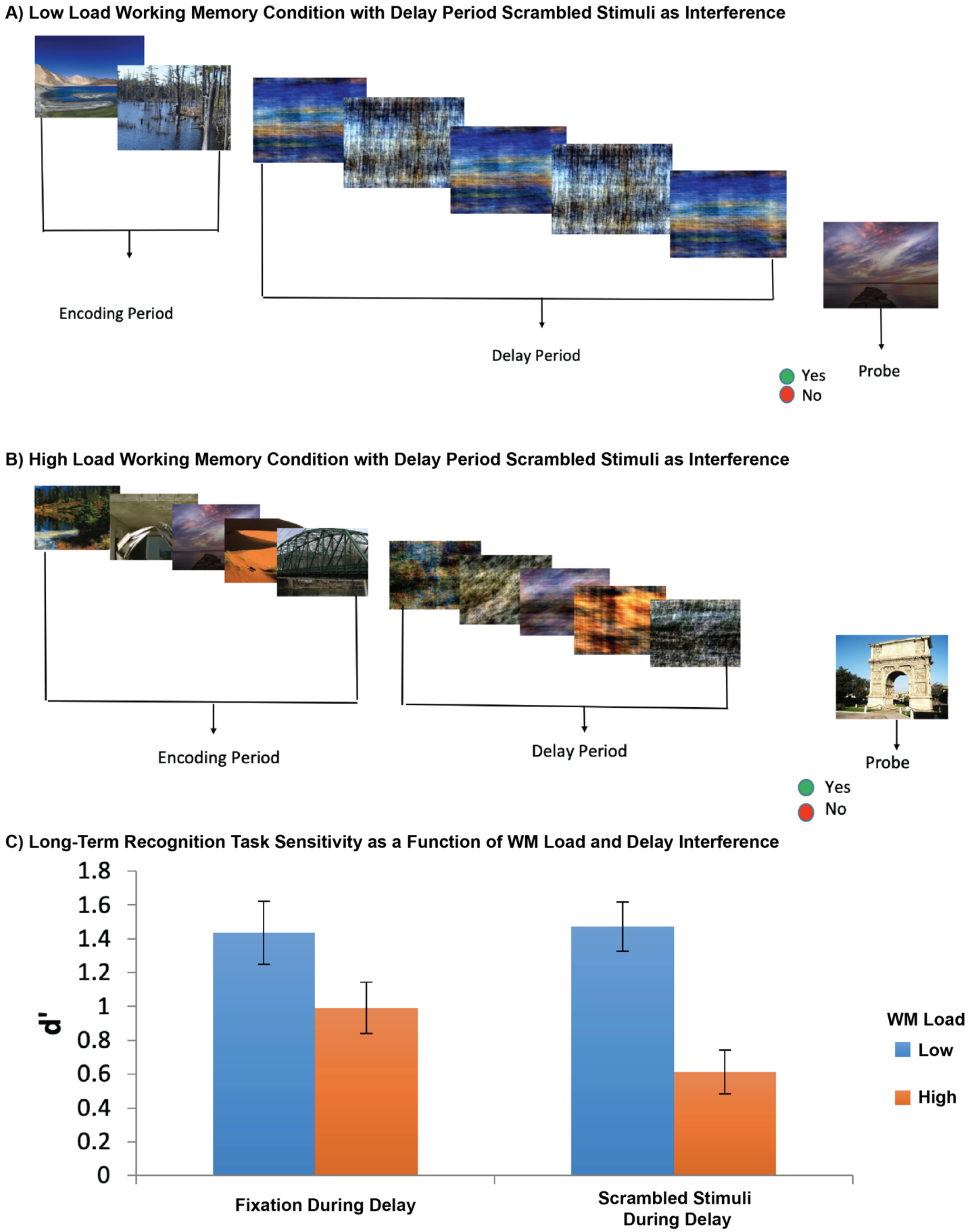


**Supplemental Figure 1. The Separate Behavioral Experiment and Results that Motivated the EEG Experiment.** Example trial for the low load WM condition with scrambled stimuli during presented during the delay period (a). Two scene stimuli were presented during the encoding period, followed by phase scrambled versions of the scenes, and then a probe stimulus to which the subject signaled whether they did or did not remember it from the set of scenes presented during encoding. Example trial of the high load WM condition with scrambled stimuli during the delay (b). In the high load condition, five images were presented during the encoding period, followed by phase scrambled versions of the scenes, and then a probe to which the subject answered whether they remembered it from the set of scenes presented at encoding. Barplot of d-prime (d') sensitivity during the long-term recognition task (c). The barplot displays the mean (±SE) sensitivity as a function of WM load (low vs high) and condition (scrambled vs. fixation). A post-hoc test revealed a significant difference between the high-load fixation and high-load scrambled delay period condition ($p = 0.002$). Sensitivity was significantly worse on the long-term recognition task for stimuli that were presented during the high-load WM condition with scrambled stimuli during the delay (mean d' = 0.41, SD = 0.18) compared to simple fixation during the delay without scrambled stimuli (mean d' = 0.79, SD = 0.43).

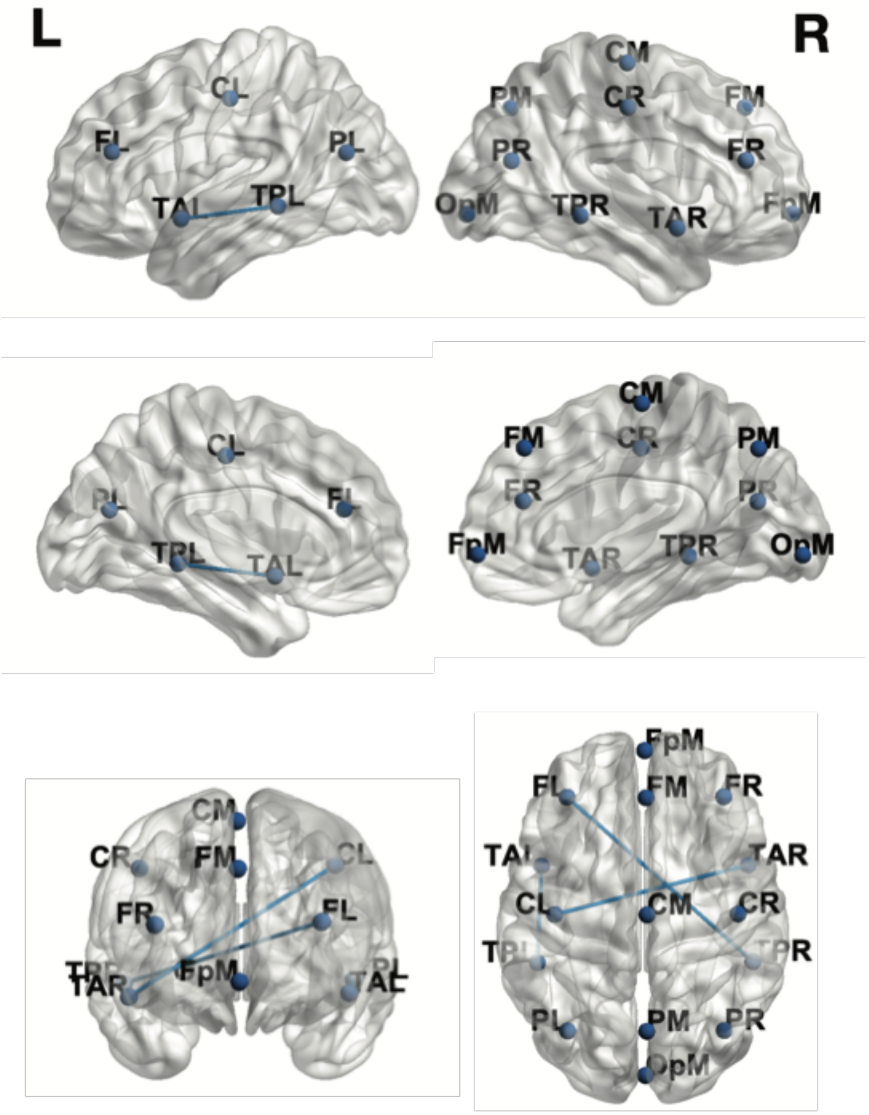


**Supplemental Figure 2. 3D Glass Brain Showing Source Analysis Brain Regions Montage**. Coordinates at each brain region indicate the center of the modeled source displayed on a glass brain (see *Supplemental Table 1*). Each brain region is represented by a Node (F=Frontal, T=Temporal, P=Parietal, O=Occipital) and lines represent functional connections that were significantly greater for the low- compared to high-load condition in the planned analyses (see *Results and Figure 6*). 3D glass brain views were generated using BrainNet Viewer (Xia, Wang, & He, 2013). The views displayed are (from left to right): [top row] left lateral, top-down, right lateral; [middle row] left medial, inferior; [bottom row] anterior, posterior.

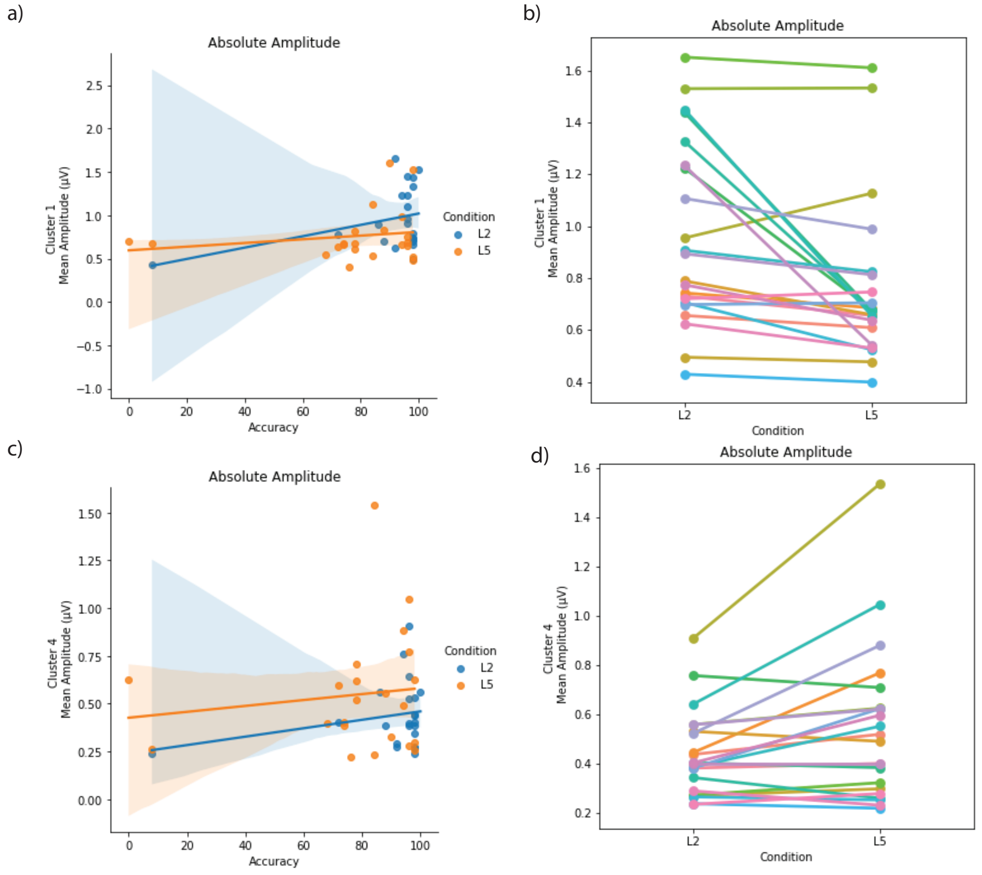


**Supplemental Figure 3. Correlations Between Clusters of Delay Activity and WM Performance**. Correlations and pointplots for Cluster 1 and Cluster 4 (*Table 1*) of the absolute amplitude comparison between the low- (L2) and high-load (L5) conditions with all trials included. Clusters were selected because they represent the largest significant clusters for the low-load greater than the high-load (Cluster 1) and the high-load greater than the low-load (Cluster 4) comparisons. a) Mean absolute amplitude for Cluster 1 for each subject (y-axis) plotted against performance as measured by accuracy (x-axis). Blue dots and lines represent the low-load condition and orange dots and lines represent the high-load condition. Regression line with bootstrapped CI interval (translucent band) plotted for each correlation. b) Mean absolute amplitude for Cluster 1 for each subject for the low- and high-load conditions. Single subject is connected by a line. Each subject is represented by a different color. c) Mean absolute amplitude for Cluster 4 for each subject (y-axis) plotted against performance as measured by accuracy (x-axis). Blue dots and lines represent the low-load condition and orange dots and lines represent the high-load condition. Regression line with bootstrapped CI interval (translucent band) plotted for each correlation. d) Mean absolute amplitude for Cluster 4 for each subject for the low- and high-load condition. A single subject is represented by a connected line and each subject is represented by a different color.

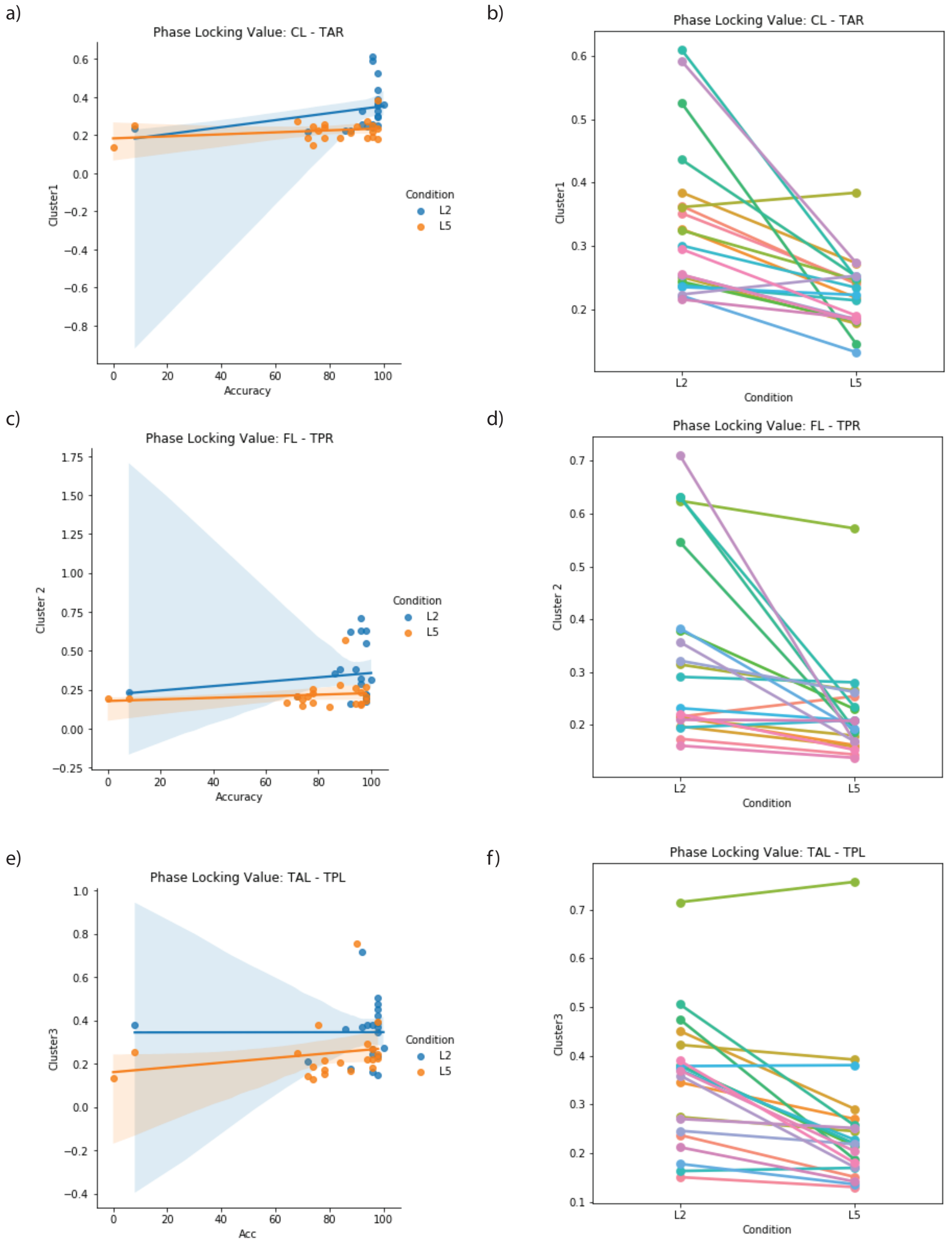


**Supplemental Figure 4. Correlations Between Clusters of Connectivity and WM Performance**. Correlations and pointplots for three significant clusters (*Table 2*) found in the phase-locking value (PLV) comparison between the low- (L2) and high-load (L5) conditions with all trials included. a) Mean PLV for Cluster 1 (CL – TAR) for each subject (y-axis) plotted against performance as measured by accuracy (x-axis). Blue dots and lines represent the low-load condition and orange dots and lines represent the high-load condition. Regression line with bootstrapped CI interval (translucent band) plotted for each correlation. b) Mean PLV for Cluster 1 for each subject for the low- and high-load condition. Each subject is connected by a line and is represented by a different color. c) Mean PLV for Cluster 2 (FL – TPR) for each subject (y-axis)

plotted against performance as measured by accuracy (x-axis). Blue dots and lines represent the low-load condition and orange dots and lines represent the high-load condition. Regression line with bootstrapped CI interval (translucent band) plotted for each correlation. d) Mean PLV for Cluster 2 for each subject for the low- and high-load condition. Each subject is connected by a line and is represented by a different color. e) Mean PLV for Cluster 3 (TAL – TPL) for each subject (y-axis) plotted against performance as measured by accuracy (x-axis). Blue dots and lines represent the low-load condition and orange dots and lines represent the high-load condition. Regression line with bootstrapped CI interval (translucent band) plotted for each correlation. f) Mean PLV for Cluster 3 for each subject for the low- and high-load condition. Each subject is connected by a line and is represented by a different color.

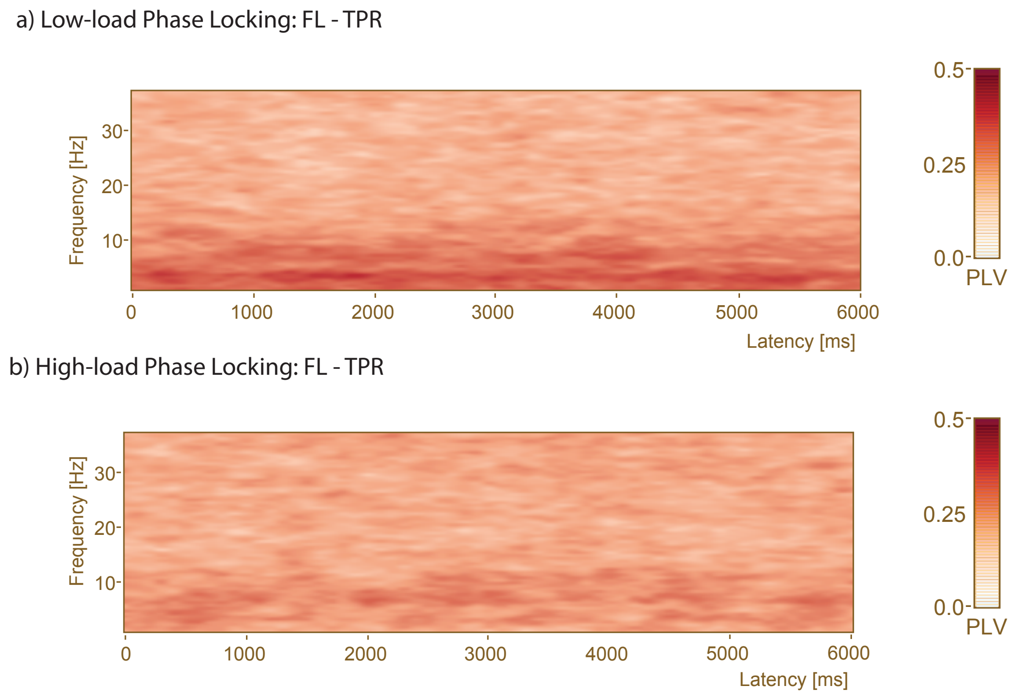


**Supplemental Figure 5. Increased Phase Locking Between FL - TPR at Low- and High-Load**. The x-axis is delay period time and the y-axis is frequency (4 – 40 Hz). The phase-locking value (PLV) plot shows phase-locking with a time-frequency bin of 0.5 Hz with 100 msec steps and on a scale from 0 (no phase-locking) to 1 (complete phase-locking). a) Phase-locking plot for the low-load condition between FL – TPR during the delay period (time: 0 – 6000 msec). b) Phase-locking plot for the high-load condition between FL – TPR during the delay period (time: 0 – 6000 msec). The PLV analysis including all trials showed a significant difference in PLV between conditions, with greater PLV during low-load throughout the entire delay period in the theta and alpha bands (see *Figure 5a*). Review of the raw PLV during low-load reveals greater phase-locking in the same frequency bands (i.e., deeper red), but the difference was not statistically significant.

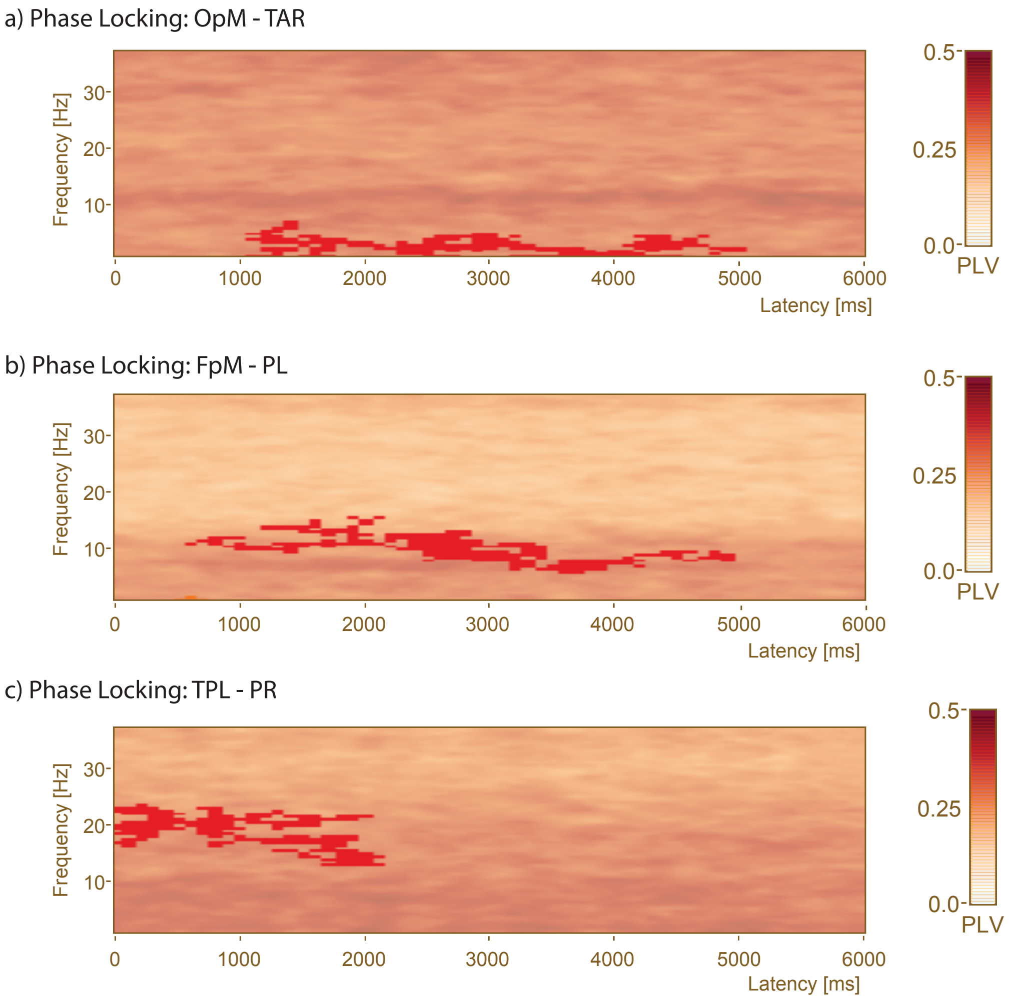


**Supplemental Figure 6. Low- and High-Load Comparison of Phase Locking with Correct Trials Only Reveals Three Additional Significant Connectivity Clusters.** The x-axis is delay period time and the y-axis is frequency (4 – 40 Hz). The phase-locking value (PLV) plot shows phase-locking with a time-frequency bin of 0.5 Hz with 100 msec steps and on a scale from 0 (no phase-locking) to 1 (complete phase-locking). The PLV analysis using only delay periods on correct response trials revealed three new clusters of different PLV. The raw PLV data is displayed with a red mask superimposed that represents significant clusters that are greater for the low-load condition. a) New cluster of different PLV for FpM – PL (time: 600-4900 msec) in the alpha and low beta band. b) New cluster of different PLV for OpM – TAR (time: 1100-5000 msec) in the theta and alpha band. c) New cluster of different PLV for TPL – PR (time: 0-2100 msec) in the low and upper beta band.

| | | MNI Coordinates (mm) | | | Brodmann Area (BA) |
|---|---|---|---|---|---|
| Hemisphere | Brain Region | X | Y | Z | |
| Midline | FpM | 0.9 | 60.6 | -5.3 | BA 10 (inferior/anterior) |
| | FM | 1.6 | 38.5 | 44.6 | |
| | CM | 2 | -16.5 | 65.4 | BA 6 |
| | PM | 1.9 | -71 | 44.3 | BA 7 |
| | OpM | 1.3 | -91.6 | -5.6 | Visual Association |
| Left Hemisphere | FL | -35.3 | 38.8 | 21.1 | Border BA 9/10 |
| | TAL | -47.3 | 6.5 | -10.3 | BA 22 |
| | CL | -41.1 | -16.4 | 46.1 | Border primary motor/sensorimotor |
| | TPL | -49.2 | -38.8 | -4.4 | BA 21 |
| | PL | -35 | -70.8 | 20.8 | Border BA 19/39 |
| Right Hemisphere | FR | 37.8 | 39 | 19.8 | Border BA 9/10 |
| | TAR | 49.2 | 6.8 | -12 | BA 22 |
| | CR | 44.5 | -16.1 | 44.6 | Border primary motor/sensorimotor |
| | TPR | 51.5 | -38.4 | -6.1 | BA 21 |
| | PR | 38.1 | -70.6 | 19.6 | Border BA 19/39 |

**Supplemental Table 1.** The MNI coordinates for the source analysis brain regions montage that displayed on the 3D glass brain views (see *Supplemental Figure 1*). The coordinates were generated from source space in BESA Research. Each coordinate represents the center of the source in a given brain region (F= Frontal, T= Temporal, P= Parietal, O=Occipital). When available the approximate corresponding Brodmann's Area (BA) is listed. These corresponding areas were identified using an MNI-Brodmann conversion tool (Lacadie, Fulbright, Arora, Constable, & Papademetris, 2008).

| Cluster ID | p-value | Cluster value | Mean for L2 Abs Amp | Mean for L5 Abs Amp | Start Time | End Time | Start Frequency | End Frequency | Electrodes In Cluster |
|---|---|---|---|---|---|---|---|---|---|
| **Cluster 1** | 0.002 | 16704.6 | 0.955287 | 0.760351 | 0 | 6000 | 4 | 40 | F3'_avr, F4'_avr, C3'_avr, C4'_avr, P3'_avr, P4'_avr, O1'_avr, O2'_avr, F7'_avr, F8'_avr, T7'_avr, T8'_avr, P7'_avr, P8'_avr, Fz'_avr, Cz'_avr, Pz'_avr, Oz'_avr, FC1'_avr, FC2'_avr, CP1'_avr, CP2'_avr, FC5'_avr, FC6'_avr, CP5'_avr, CP6'_avr, POz'_avr |
| **Cluster 2** | 0.012 | 2358.94 | 1.28309 | 0.904188 | 0 | 6000 | 4 | 18.5 | FP1'_avr |
| **Cluster 3** | 0.024 | 1183.24 | 1.08534 | 0.816534 | 0 | 6000 | 4 | 16.5 | FP2'_avr |
| **Cluster 4** | 0.025 | -1032.25 | 0.369919 | 0.464164 | 0 | 3500 | 30 | 37 | C4'_avr, P4'_avr, CP2'_avr, FC6'_avr, CP6'_avr |
| **Cluster 5** | 0.041 | 563.23 | 0.78447 | 0.656996 | 2200 | 4400 | 14.5 | 19.5 | F3'_avr, C3'_avr, P4'_avr, Fz'_avr, Pz'_avr, FC1'_avr, CP1'_avr, CP2'_avr, POz'_avr |
| **Cluster 6** | 0.038 | -484.496 | 0.349476 | 0.433934 | 3700 | 6000 | 30.5 | 37 | C4'_avr, P4'_avr, CP2'_avr, FC6'_avr, CP6'_avr |

**Supplemental Table 2.** Significant clusters of absolute amplitude difference between the low- (L2) and high-load (L5) conditions during the delay period (Time: 0 to 6000 msec) for all sensors. This table contains the results for the analysis of absolute amplitude between the conditions using trials with correct only responses.

| Cluster ID | p-value | Cluster value | Mean for L2 | Mean for L5 | Start Time | End Time | PLV Regions |
|---|---|---|---|---|---|---|---|
| **Cluster 1** | 0.009 | 834.882 | 0.356729 | 0.232638 | 700 | 6000 | CL-TAR |
| **Cluster 2** | 0.009 | 834.882 | 0.356729 | 0.232638 | 700 | 6000 | TAR-CL |
| **Cluster 3** | 0.036 | 519.548 | 0.377551 | 0.2672 | 600 | 4900 | PL-FpM |
| **Cluster 4** | 0.036 | 519.548 | 0.377551 | 0.2672 | 600 | 4900 | FpM-PL |
| **Cluster 5** | 0.05 | 448.305 | 0.413132 | 0.291091 | 1100 | 5000 | OpM-TAR |
| **Cluster 6** | 0.05 | 448.305 | 0.413132 | 0.291091 | 1100 | 5000 | TAR-OpM |
| **Cluster 7** | 0.051 | 446.794 | 0.38855 | 0.282937 | 0 | 2100 | TPL-PR |
| **Cluster 8** | 0.051 | 446.794 | 0.38855 | 0.282937 | 0 | 2100 | PR-TPL |
| **Cluster 9** | 0.086 | 359.989 | 0.373 | 0.263246 | 0 | 2300 | TAL-TPL |
| **Cluster 10** | 0.086 | 359.989 | 0.373 | 0.263246 | 0 | 2300 | TPL-TAL |
| **Cluster 11** | 0.1 | 336.787 | 0.364389 | 0.258161 | 3200 | 5700 | TAL-TPL |
| **Cluster 12** | 0.1 | 336.787 | 0.364389 | 0.258161 | 3200 | 5700 | TPL-TAL |

**Supplemental Table 3:** Significant clusters of phase-locking value (PLV) between 15 brain regions during the delay period (Time: 0 to 6000 msec). Each PLV connection between the low- (L2) and high-load (L5) conditions has two significant clusters associated with it to represent the opposite connection (e.g., PL – FL and FL – PL). The PLV results do not provide information about the directionality. This table contains the results for the PLV analysis using trials with correct only responses.
